# Modelling structural constraints on protein evolution via side-chain conformational states

**DOI:** 10.1101/530634

**Authors:** Umberto Perron, Alexey M. Kozlov, Alexandros Stamatakis, Nick Goldman, Iain H. Moal

## Abstract

Few models of sequence evolution incorporate parameters describing protein structure, despite its high conservation, essential functional role and increasing availability. We present a structurally-aware empirical substitution model for amino acid sequence evolution in which proteins are expressed using an expanded alphabet that relays both amino acid identity and structural information. Each character specifies an amino acid as well a rotamer state: the discrete geometric pattern of permitted side-chain atomic positions. By assigning rotamer states in 251,194 protein structures and identifying 4,508,390 substitutions between closely related sequences, we generate a 55-state model that shows that the evolutionary properties of amino acids depend strongly upon side-chain geometry. The model performs as well as or better than traditional 20-state models for divergence time estimation, tree inference and ancestral state reconstruction. We conclude that the concomitant evolution of sequence and structure is a valuable source of phylogenetic information.

## Introduction

The development of evolutionary models is a prerequisite (albeit sometimes an implicit one) for many common bioinformatics tasks such as recognition of homologous sequences, phylogenetic tree estimation, evolutionary hypothesis testing and protein structure prediction (Huelsenbeck and Rannala 1997, Felsenstein 2004, Koonin 2005, Ginalski 2006). Because of this, the development and improvement of model-based approaches to studying protein evolution is an area of research where advances have wide-spread benefits. Furthermore, high-resolution structural information from a variety of techniques is now available for large numbers of proteins and molecular assemblies (Milne *et al.* 2013, Carroni and Saibil 2016, Venien-Bryan *et al.* 2017), improving our understanding of how protein folding and residue function change over time.

### Empirical models of amino acid replacement

When studying the evolution of amino acid sequences, substitutions are usually described using a continuous-time Markov model with the 20 amino acids as the states of the chain (Liò *et al.* 1998, Felsenstein 2004, Thorne and Goldman 2007, Perron *et al.* ressin press). Models belonging to the empirical class are built by analysing large quantities of sequence data (typically hundreds of protein alignments) and estimating relative substitution rates between all state (amino acid) pairs under a time-reversible model. Empirical models are typically assumed to be applicable to broad classes of proteins with little further parameter optimization aside from techniques that match amino acid frequencies to what is observed in a specific dataset under study and allow for rate heterogeneity amongst sequence sites (Yang 1993).

The first empirical amino acid substitution model was introduced by Dayhoff and colleagues in 1966 (Eck and Dayhoff 1966), and updated regularly until 1978 (Dayhoff *et al.* 1978). They compiled protein sequence alignments and tabulated amino acid substitutions along branches on the phylogenetic trees; from these data, an instantaneous rate matrix *Q* defining a continuous-time Markov model can be constructed (Kosiol and Goldman, 2005). Since then, a number of alternative *Q*-matrices for amino acid substitution have been developed using similar approaches but taking advantage of more powerful model estimation techniques, and larger or more specific datasets. Examples include MTMAM (Yang *et al.*, 1998) for mammalian mitochondrial proteins and GPCRtm for the transmembrane region of GPCRs (Rios *et al.*, 2015); WAG (Whelan and Goldman, 2001) and LG (Le and Gascuel, 2008) that were derived from larger, diverse databases of sequence alignments; and LG4X (Le *et al.*, 2012) that aims to capture varying evolutionary dynamics at different sequence sites.

### Role of structural information in understanding protein evolution

Although considering particular proteins’ distinct amino acid compositions and among-site rate variation improves a model’s statistical fit to the data indicating a better description of evolutionary patterns, it seems clear, from considering protein structure and function, that at least some variability in the evolutionary process will be associated with the structural environment of a site (Thorne and Goldman 2007, Perron *et al.* ressin press). For example, solvent-exposed residues evolve more rapidly than those buried in the protein interior (∼×2) and exhibit different amino acid frequencies and substitution patterns, due to less steric constraint and the need to interact favourably with water. Similarly, secondary structure also influences substitutions, for instance disfavouring amino acids in *α*-helices that are incompatible with the canonical *α*-helical conformation due to disrupting backbone geometry or steric clashes arising from branching at the C^*β*^ atom. Models that account for some of these differences, for instance by using a separate 20-state model for different structural environments, have been shown to improve over those based on sequence alone (Overington *et al.* 1992, Goldman *et al.* 1998, Liò *et al.* 1998).

The tertiary and quaternary structures of proteins provide further constraints to evolution, in the form of natural selection operating on the specific inter-residue interactions that stabilise the fold of the protein, as well as the need to avoid misfolding and aggregation (Overington *et al.* 1990, Shakhnovich *et al.* 1996). While attempts to model some of these factors have been made (Robinson *et al.* 2003, Bastolla *et al.* 2003, 2006, Rodrigue *et al.* 2005, 2006, Arenas *et al.* 2017), this has proven difficult due to the challenging computational requirements of models that allow evolution at one position to be dependent upon the sequence at other positions. In addition to the difficulties that arise when site independence can no longer be assumed, these models are considerably more complex and require further assumptions such as (1) a constant tertiary structure, (2) approximate functions to map sequence to stability or misfolding propensity and (3) additional approximate functions to map stability or misfolding propensity to rate effects. They are computationally demanding, difficult to use in an inference setting, and are not, unlike our model, readily integrated into commonly used software.

### An evolutionary model based on side-chain rotamer states

In this paper, we present an evolutionary model that introduces structural information by accounting for the conformational state of each residue based on the atomic positions of its side-chain. Specifically, we split the traditional 20 amino acid states into discrete sub-states based on the *χ*_1_ rotamer (short for ‘rotational isomer’) configuration of their side-chains as defined in the Dunbrack rotamer library (Shapovalov and Dunbrack, 2011). In this classification, each residue can adopt one of (typically) three discrete configurations (Fig. 1, Sup. Fig. 1) defined by the dihedral angle between the first two covalently linked carbons in the side-chain (*C*^*α*^ and *C*^*β*^). These three stable rotamer configurations correspond to specific *χ*_1_ dihedral angle values (∼ 60°, ∼-180°and ∼-60°) consistently across all residues; this means that residues sharing the same rotamer configuration (e.g. PHE1 and TRP1 in Fig. 1) have side-chains that are similarly oriented with respect to the backbone. The adoption of one rotamer configuration over another is determined by their relative stability, a combination of the intrinsic stability of that state, local factors such as the backbone geometry and the position of atoms further along the side-chain, and the forces applied by the surrounding residues and the requirement to pack alongside them. Thus, they convey information about both the local structure as well as the interactions of the residue within the fold as a whole. As these states are discrete and finite, and each residue in a protein structure adopts exactly one *χ*_1_ configuration, they can be readily incorporated into an expanded alphabet of amino acid states, maintaining the usual assumption of sitewise independence. This produces a scalable model that can be used in the same way as a traditional 20-state substitution model.

**FIG. 1.**
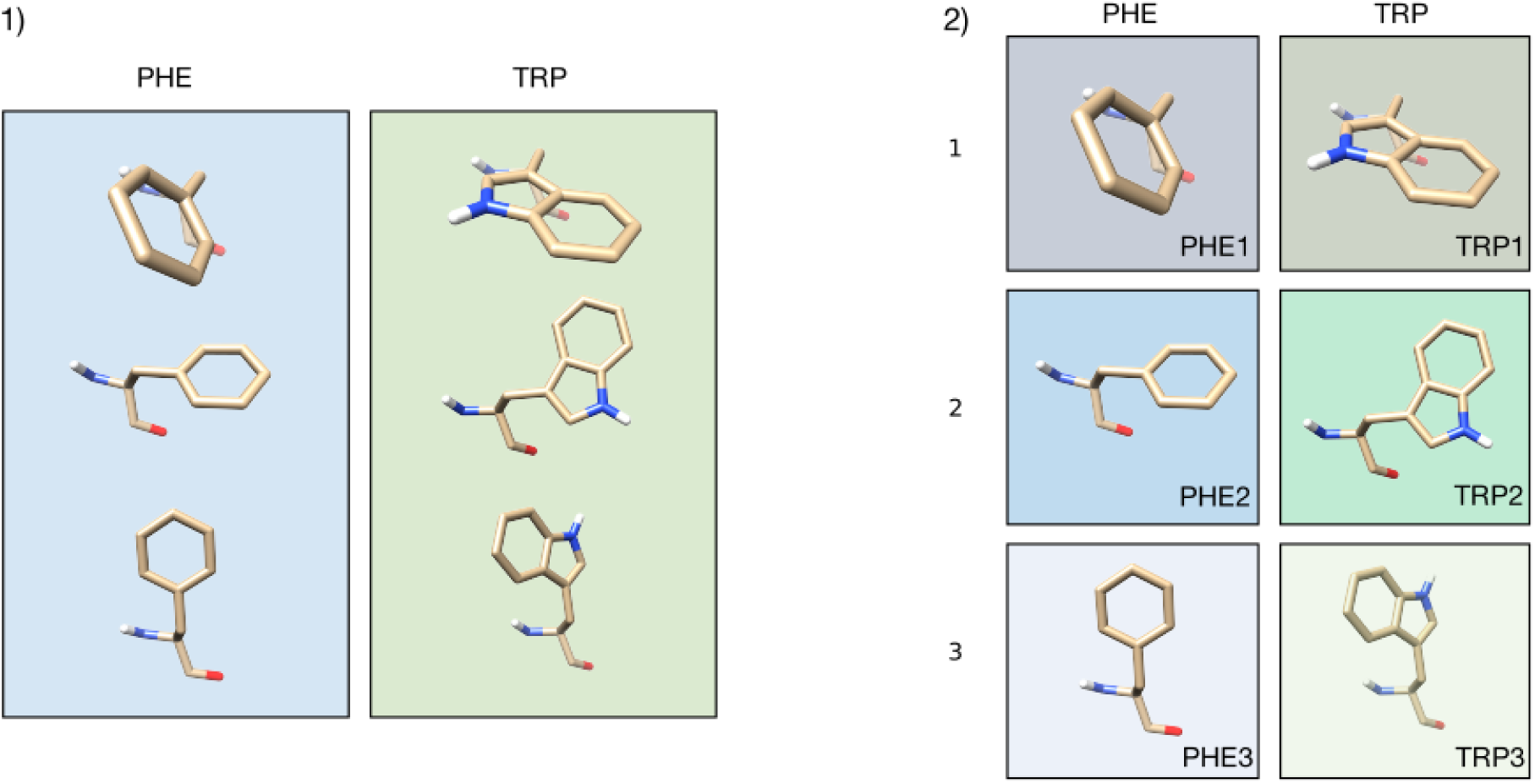
Illustration of the rotamer configurations of phenylalanine (PHE) and tryptophan (TRP). **(a)** In traditional amino acid replacement models their distinct *χ*_1_ rotamer configurations are merged into a single amino acid state. **(b)** In our model these states are split into three *χ*_1_ configuration-specific states (1, 2, and 3) defined, as in the Dunbrack rotamer library, by the dihedral angle between the first two covalently linked carbons in the side-chain (*C*^*α*^ and *C*^*β*^; see also Sup. Fig. 1).

By compiling a large set of homologous sequences for which structural data are available, we develop a structurally-aware “Dayhoff-like” substitution model based on an instantaneous rate matrix that uses an expanded state set composed of 55 states, each of which corresponds to the combination of a residue and its *χ*_1_ configuration (Table 1). Almost all amino acids show a significant, and often strong, conformational dependence in their substitution patterns, indicating that an amino acid can behave as a distinct entity depending on the orientation of its side-chain. Thus, our 55-state model (denoted ‘RAM55’, for Rotamer-Aware Model) introduces valuable, biochemically plausible, structural information while retaining a classic architecture that can be readily implemented in widely-used phylogenetic inference software such as RAxML-NG (Stamatakis 2014, Kozlov *et al.* 2018). This model improves our understanding of the relationships between protein sequence, structure and evolution.

**Table 1.** Rotamer configuration states. Left: 20 traditional states correspond to the 20 amino acids. Right: Our 55-member expanded state set describes both the amino acid and *χ*_1_ rotamer configuration for each constituent residue of a protein. Most amino acids have three possible *χ*_1_ configurations corresponding to specific *χ*_1_ dihedral angle values (∼ 60°, ∼ - 180°and ∼ - 60°) (see Sup. Fig. 1). Alanine (ALA) and glycine (GLY) have no side-chain and therefore no *χ*_1_ configuration, while proline (PRO) only has two stable *χ*_1_ configurations (∼-27°, ∼ - 25°) because of steric requirements of its pyrrolidine ring.

| Traditional state | Expanded states |  |  |
| --- | --- | --- | --- |
| ALA | ALA |  |  |
| ARG | ARG1 | ARG2 | ARG3 |
| ASN | ASN1 | ASN2 | ASN3 |
| ASP | ASP1 | ASP2 | ASP3 |
| CYS | CYS1 | CYS2 | CYS3 |
| GLN | GLN1 | GLN2 | GLN3 |
| GLU | GLU1 | GLU2 | GLU3 |
| GLY | GLY |  |  |
| HIS | HIS1 | HIS2 | HIS3 |
| ILE | ILE1 | ILE2 | ILE3 |
| LEU | LEU1 | LEU2 | LEU3 |
| LYS | LYS1 | LYS2 | LYS3 |
| MET | MET1 | MET2 | MET3 |
| PHE | PHE1 | PHE2 | PHE3 |
| PRO | PRO1 | PRO2 |  |
| SER | SER1 | SER2 | SER3 |
| THR | THR1 | THR2 | THR3 |
| TRP | TRP1 | TRP2 | TRP3 |
| TYR | TYR1 | TYR2 | TYR3 |
| VAL | VAL1 | VAL2 | VAL3 |

We further show that RAM55 results in a detectable improvement in model fit on simulated data, and on a number of diverse empirical datasets. It produces reliable tree topology and sequence divergence estimates. In addition, the RAM55 model also allows structurally-aware reconstruction of both ancestral rotamer and amino acid states. This is of relevance to ancestral sequence reconstruction/resurrection used, for example, to investigate how the physical properties of proteins shaped their evolutionary process (e.g. Harms and Thornton 2013; Wheeler *et al.* 2016). We show that RAM55 can accurately reconstruct ancestral rotamer states from descendant protein sequences of known structure; it is also able to reconstruct ancestral amino acid states as well as or better than traditional 20-state models. Reconstructed rotamer states could help in resurrecting ancestral proteins by providing insight into their secondary structures as certain rotamer configurations are only allowed within a specific backbone geometry (Dunbrack and Cohen 1997, Lovell *et al.* 2000, Dunbrack 2002).

## Results and Discussion

### Rotamer state exchange rates

We first investigate how rotamer states exchange over evolutionarily-relevant time-spans by computing a replacement rate matrix derived from counting changes in homologous sites of proteins of known structure (see Materials and Methods: *Rotamer assignment and sequence alignments* and *Tabulating substitution counts*). Figure 2 shows these exchangeabilities in heat-map form for our 55-state model (RAM55), and for a 20-state ‘rotamer-unaware’ empirical model (RUM20) we estimated from the same dataset for comparison purposes. Our exchange rates show evidence of *χ*_1_ configuration conservation, whereby the *χ*_1_ configuration (*R*) is frequently conserved when the identity of the amino acid (*A*) changes (i.e. (*A,R*) ↔ (*A*′*,R*) with *A*′ ≠ *A*). This is visible in Fig. 2 where higher exchange rates are observed on the diagonal of many of the 3×3 sub-matrices (corresponding to changes in amino acid only) compared to the off-diagonal elements (changes in amino acid and rotamer configuration). This is particularly true of interchanges between biochemically similar amino acids: sub-matrices corresponding to aromatic-aromatic exchanges for example all have very distinct diagonal patterns, as do the exchanges between aspartic acid (ASP) and its derivative asparagine (ASN), and between serine (SER) and threonine (THR).

**FIG. 2.**
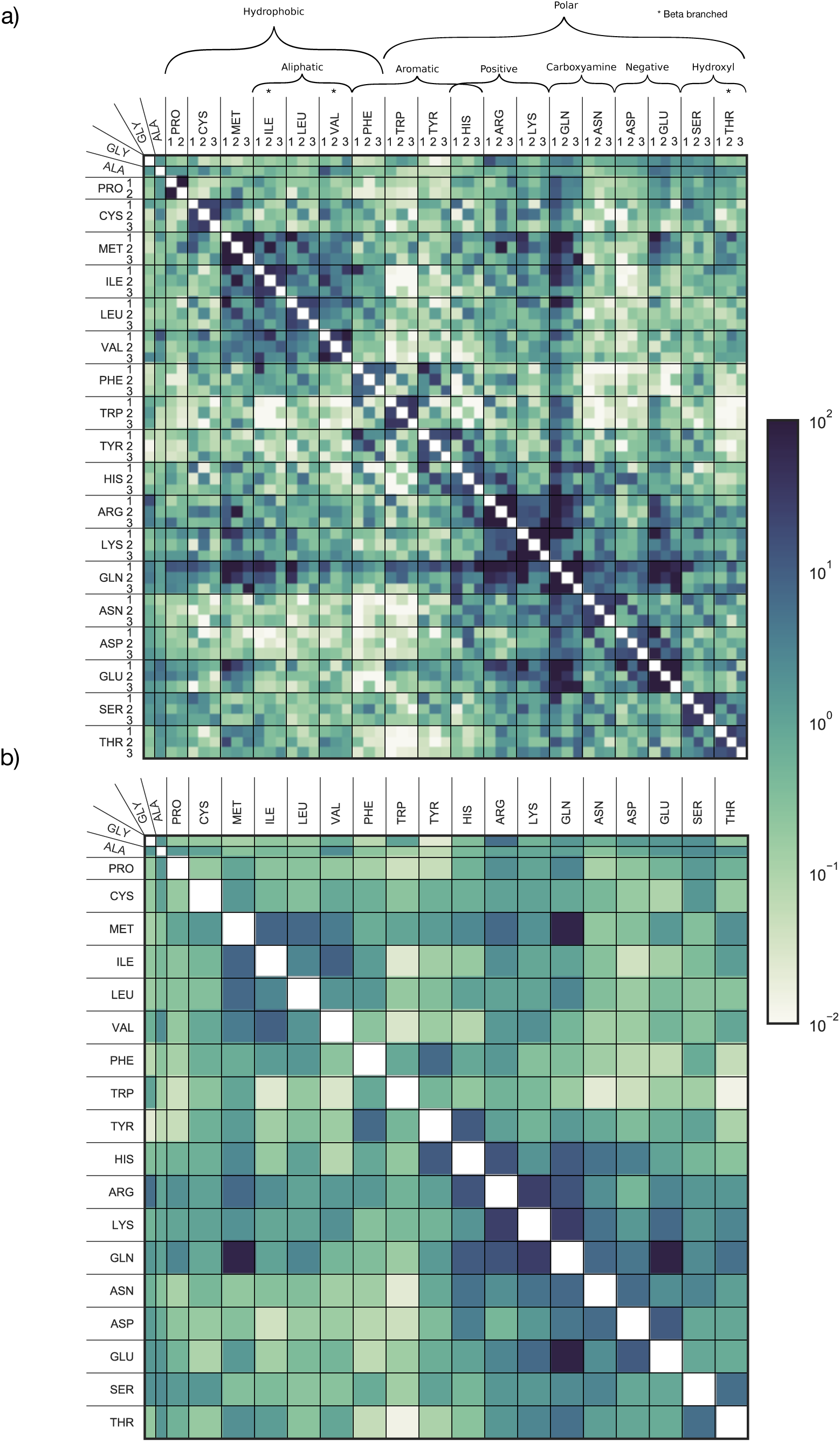
Replacement models, with and without rotamer configuration information. Exchangeabilities (*s*_*(A,R)*,(*A*′,*R*′)_ and *s*_*A,A*′_) are reported in heat-map form for **(a)** our 55-state model (RAM55) and **(b)** a 20-state model (RUM20) estimated from the same dataset. Note that time-reversibility of the models means the exchangeabilities are symmetric (e.g. *s*_*(A,R)*,(*A*′,*R*′)_ = *s*_(*A*′,*R*′)*,(A,R)*_ for (*A,R*) ≠(*A*′*,R*′)).

Overall, 111 of the 136 independent 3×3 sub-matrices show significant association among the interchanging states (see *Rotamer state exchangeability analysis*). To further quantify the strength of these sub-matrix patterns we use Cramér’s 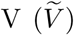, a measure of association between two categorical variables (here the *χ*_1_ configurations of amino acids *A* and *A*′). Existence of strong association does not guarantee a diagonal pattern (rotameric state conservation); we therefore also consider the diagonal ratio for each sub-matrix, indicating the tendency of rates to lie on each 3×3 sub-matrix’s diagonal. 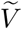 and diagonal ratio are shown in Fig. 3 for the 111 sub-matrices with significant associations.

**FIG. 3.**
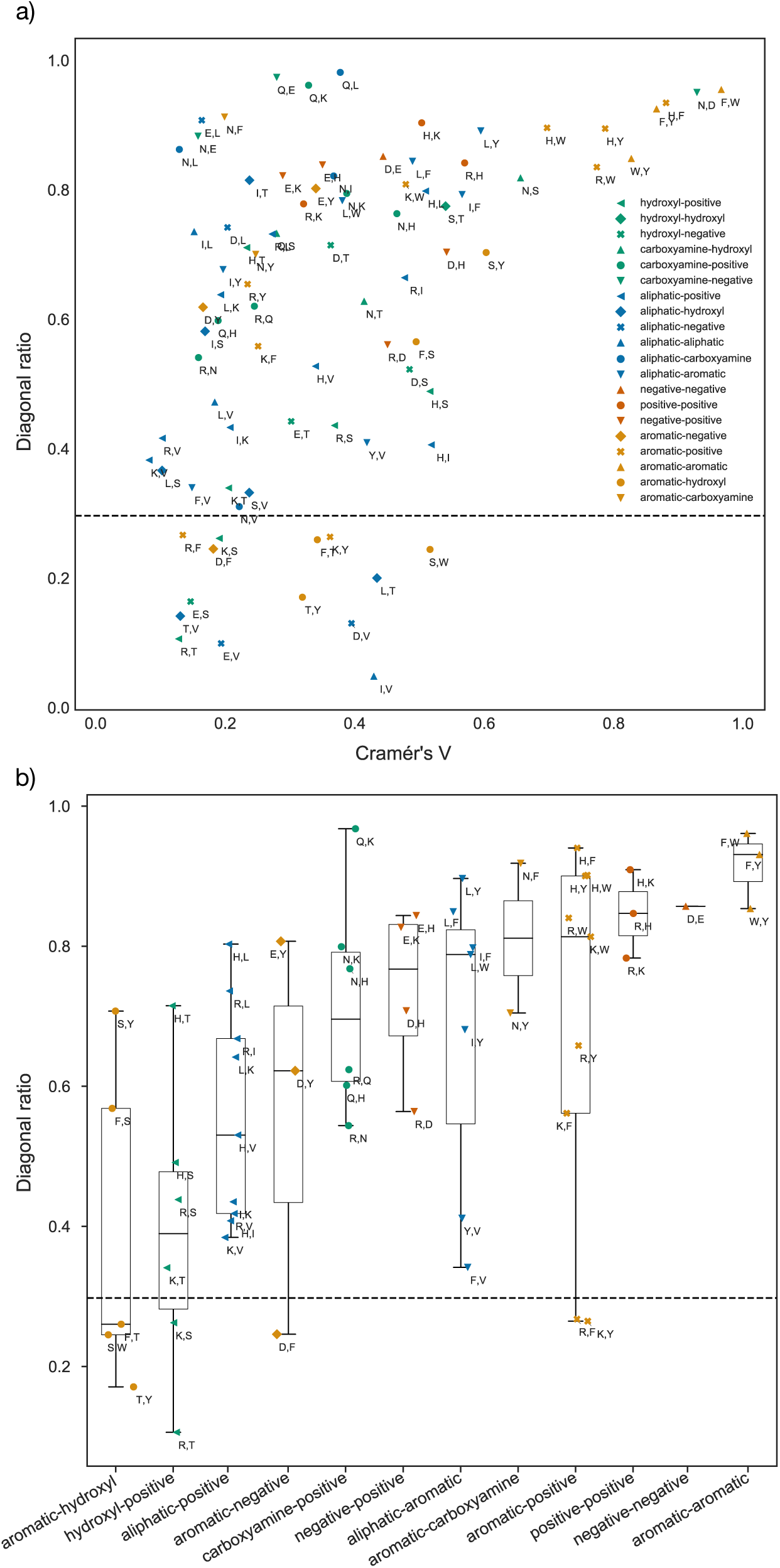
Strength of association and diagonal ratio. Plots show pairs of residues whose 3 × 3 sub-matrices within the RAM55 *Q*-matrix achieve significant *χ*^2^ statistic values. Pairs are labelled according to their component residues’ biochemical properties. **(a)** Strength of association (Cramér’s 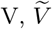) between the *χ*_1_ configurations of residues composing each pair and diagonal ratio (measuring propensity to conserve *χ*_1_ configuration). **(b)** Box-plots show diagonal ratio values and medians for exchanges between pairs of residues grouped according to biochemical similarities.

All six aromatic-aromatic sub-matrices have high 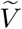 values and high diagonal ratios (Fig. 3a, upper right), indicating a strong preference for conserving side-chain orientation. This exchange pattern might be capturing the effect of local constraints on how freely a bulky aromatic side-chain can be positioned without displacing or clashing with those of neighbouring residues. A similarly strict configuration conservation can be observed for negative-negative and positive-positive exchanges; however, negative-positive exchanges have high 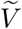 but somewhat lower diagonal ratios (Fig. 3b). These sub-matrices show significant association between specific configurations of the exchanging residues but no common pattern, possibly arising from the competing pressures to retain compatible geometries upon substitution but also to displace the charged moiety to a new location following a charge swap. It is also interesting to note that leucine has high diagonal score in exchanges with all the aromatics (aliphatic-aromatic comparisons, Fig. 3b). In contrast, isoleucine and valine, both aliphatic and *β*-branched, have lower scores and show less tendency to conserve their side-chain orientation when exchanging to aromatic residues.

In addition to the conservation of the *χ*_1_ configuration upon amino acid substitution, we also investigated the influence backbone geometry may have on the observed exchangeabilities. To do so we calculated, for each pair of rotamer states, the overlap between the bivariate joint distributions of their *ϕ* and *ψ* backbone dihedral angles. These overlap values correlate with the exchangeabilities, with a Spearman’s *ρ* of 0.29 (*p* = 1.7×10^-28^), indicating that rotamer states exhibit a weak but highly significant preference to interchange with other rotamer states that occupy similar regions of the Ramachandran plot (see *Overlap of backbone distributions*). In some cases, strong non-diagonal patterns in the exchageabilities of amino acid pairs correlate strongly with the overlap values, for instance for the exchange of threonine and leucine (*ρ* = 0.85), and indeed 76% (115 out of 153) of the 3×3 and 2×3 sub-matrices corresponding to changes in amino acid have a positive Spearman’s *ρ* between overlap and exchangeability (see Sup. Fig. 2), indicating that for most amino acid pairs there is a tendency for evolutionary exchanges to be between side-chain geometries that accommodate similar backbone geometries.

Aside from those discussed above, there are several highly significant associations that have no obvious biochemical explanation. Indeed, some cases exhibit a tendency to avoid conserving *χ*_1_ configurations during amino acid exchanges: for example, see the isoleucine-valine interchanges in Figs. 2a and 3a. Nevertheless, the strength of these associations indicates that our expanded state set incorporates valuable, biochemically plausible, structural information into the model. RAM55 (Fig. 2a) can thus be considered a ‘high-resolution’ version of RUM20 (Fig. 2b) generated from the same dataset. As we show in subsequent sections this provides additional inferential power from the ability to distinguish states and state-interchanges according to *χ*_1_ configuration.

Due to RAM55’s expanded state-space, the probability of observing any amino acid, given the initial state and a divergence time *t*, is different in the 55-state model than it is in the 20-states model. For instance, a histidine residue is more likely to be substituted with an asparigine when *χ*_1_ ≈ 60° than when in one of the other *χ*_1_ configurations. In the RUM20 model the three *χ*_1_ configurations are merged, and thus the amino acid probability distribution at time *t* corresponds to the weighted average of the three rotamer states. Thus, for each rotamer state, there is a divergence between the probability distributions of the amino acids states at time *t* using the RAM55 model when compared to that when using RUM20. Indeed, as RUM20 can be arrived at by merging states in RAM55, this divergence constitutes a loss of information regarding the amino acid probability distribution when RAM55 is approximated using RUM20. This can be quantified using the Kullback-Leibler divergence (Kullback and Leibler 1951; see *Kullback-Leibler divergence*). At *t* = 0, no loss occurs due to the amino acid sequence being fully known in both models. As *t* →∞, both models tend towards the equilibrium amino acid frequencies and the loss tends towards zero. The differences between the two models manifest in between these extremes. Figure 4 shows that average information loss for one state peaks at 0.0002 *bit* per site after 0.4 amino acid substitutions per site have occurred on average, although this can be much higher for certain rotamer states, and moreover indicates that the difference is most pronounced at the timescales at which evolutionary models are commonly applied: up to *t* = 2.5 which corresponds to ∼ 20% amino acid sequence identity.

**FIG. 4.**
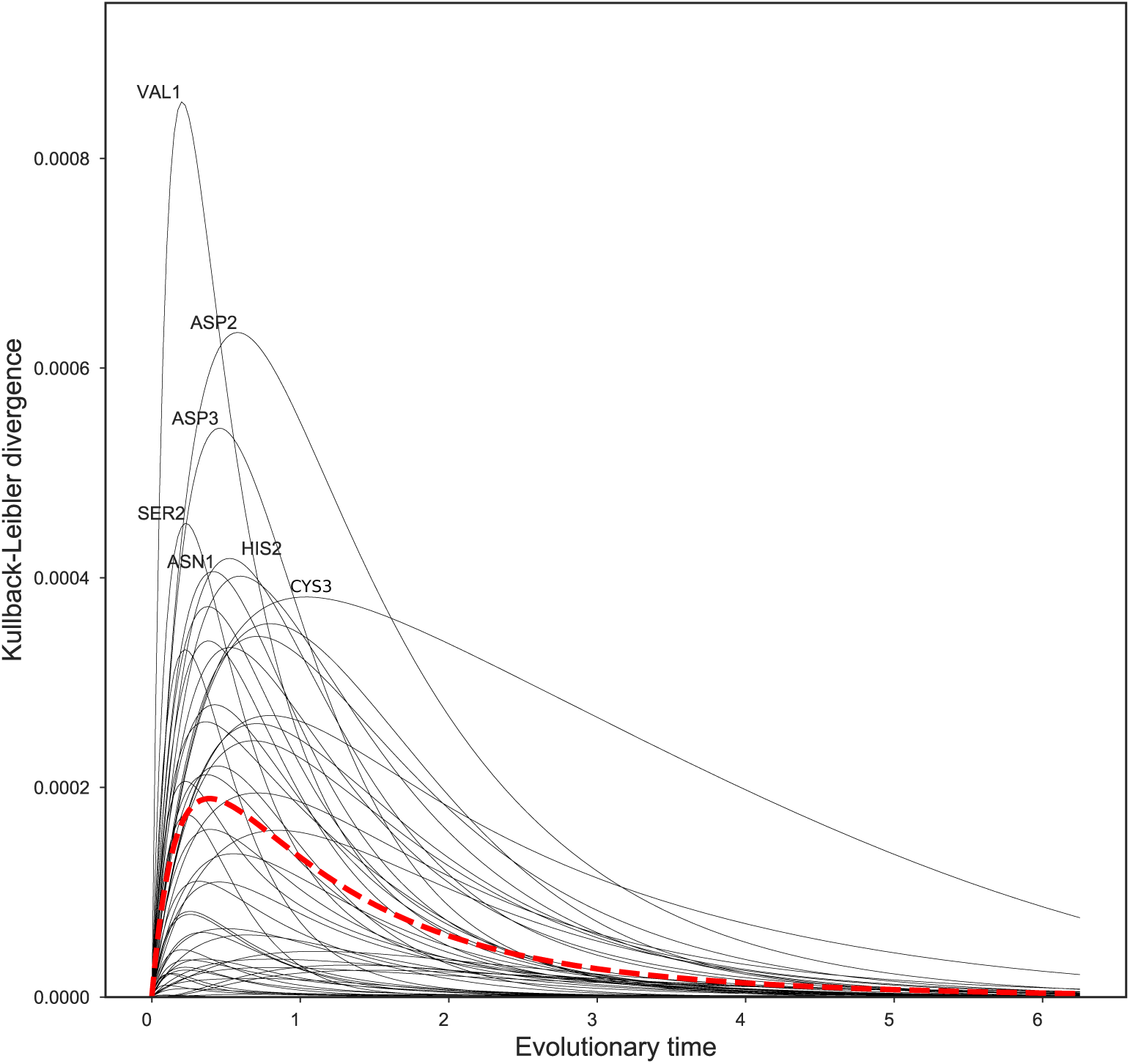
Kullback-Leibler (KL) divergence measures the amount of information lost when the structure-free RUM20 model is used to approximate the 55-state model, RAM55. KL divergence is computed for every pair of amino acid state and corresponding rotamer state as a function of evolutionary time *t* (expressed as expected number of amino acid substitutions). The overall information loss, computed by averaging over all state pairs’ KL divergences and weighted by the rotamer state equilibrium frequencies, is shown in red.

### Model benchmarking: simulation

Here we use simulated alignments to assess whether or not typical protein sequence data sets contain enough information to permit identification of the true generating model, in the case that the data were generated by the RAM55 rotamer state-aware model. We also investigate whether RAM55 affects our ability to infer phylogenies compared to models using the traditional 20-state space.

To examine our ability to detect *χ*_1_ configuration-influenced evolution, we assessed our 55-state model’s goodness-of-fit when analysing alignments simulated using the model itself. These simulations use a variety of phylogenetic trees and branch scaling factors, to allow evaluation of model detection over a wide range of realistic conditions (see *Tree generation and alignment simulation*). From these simulated alignments we then infer the corresponding phylogenies by maximum likelihood (ML) using RAM55 or other models that are widely used for phylogenetic analysis of amino acid sequences (see *Likelihood calculation and maximization over phylogenies*). Figure 5a compares AIC scores (Akaike, 1974) or state-corrected AIC scores (see *Log-likelihood comparison across models)* of the inferred tree across multiple models. RAM55 consistently shows detectably better fit for the simulated data regardless of sequence divergence. Moreover, for most branch lengths and number of taxa, RAM55 has a lower AIC score than all other models for 100% of the simulations (see Sup. Tab. 1). At the extreme of trees with large tree lengths and low taxa number, the RUM20 model occasionally has a lower AIC score.

**FIG. 5.**
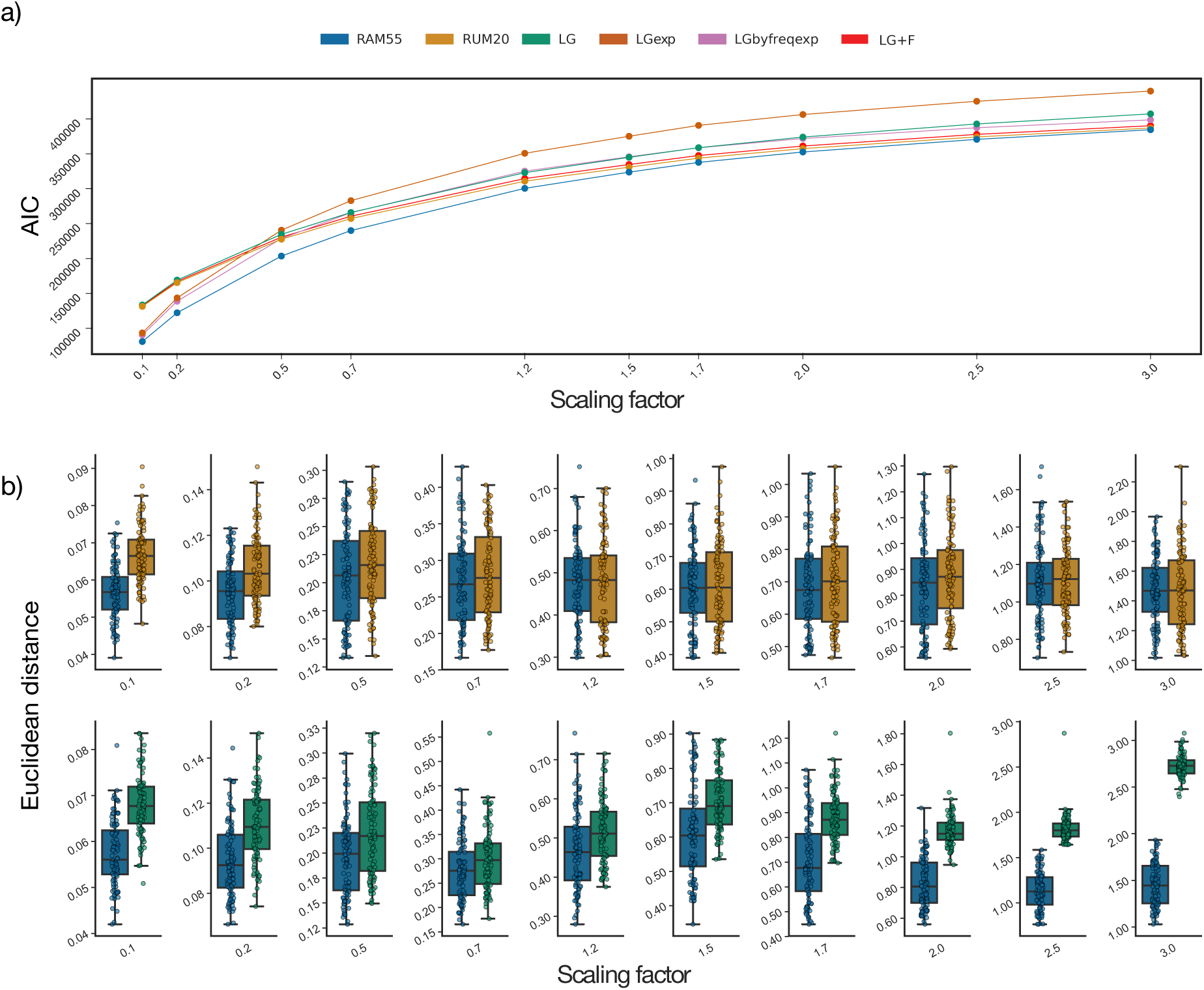
**(a)** AIC values for competing models. Each data point corresponds to the mean AIC value of trees inferred from 100 simulated alignments. **(b)** Comparison of the RAM55 model (blue bars) against LG (green; top panel) and against our RUM20 model (orange; bottom panel) in terms of Euclidean distance of their inferred trees from the reference phylogeny used to simulate the alignment. Box plots illustrate distance distribution and median (horizontal lines), scatter plot points represent individual distance values. Tree inference is performed on alignment data sets (1000 sites, 64 taxa, 100 replicates per scaling) simulated using RAM55 and a randomly-generated reference phylogeny, scaled according to the factors on the *x*-axis.

It is also interesting to note how LGexp, our version of the 20-state LG model ‘uniformly expanded’ to 55 states but incorporating no structural information (see *Log-likelihood comparison across models)*, fits the data worse than its ‘frequency-aware’ counterpart (LGbyfreq-exp) whose AIC values are comparable with those of RUM20 and LG. This illustrates how adding non-informative complexity to a 20-state model penalizes its performance, whilst being correctly informed about each rotamer state’s frequency but not specifically about its exchange rates still produces a worse fit than the full RAM55 model. These results confirm that, when the more complex RAM55 model matches the underlying process generating the input alignments, it is possible to detect an improvement in fit over simpler models.

As a further performance test, we also evaluated whether ML trees inferred under the RAM55 model are closer to the reference phylogeny used during the simulation process than those inferred with other models. For these comparisons we considered both (1) the Euclidean distance (Felsenstein 2004), a metric that accounts for both topological differences between trees and differences in branch lengths, and (2) the lengths of individual branches. Under the former measure, RAM55 infers trees that are at least as close or closer to the reference phylogenies than those inferred by amino acid replacement models such as our RUM20 model or LG. Fig. 5b compares the distributions of Euclidean distances between inferred and reference trees, estimated using the RAM55, RUM20 and LG models in simulations of 1000 sites on a 64-taxon phylogeny. Shifts towards lower values for RAM55 indicate greater accuracy of trees inferred using this model. Similar results are obtained for other alignment lengths and numbers of taxa in model phylogenies, as well as when simulating over a larger, empirical tree. (see Sup. Fig. 3–5).

Considering individual branches, RAM55 tends to more accurately estimate the correct evolutionary distance between sequences regardless of tree size (number of taxa), length of the examined branch or branch positioning in the tree. Fig. 6 — highlighting one internal branch for each of four topologies — illustrates this with branch length estimates from RAM55 having smaller variances and medians nearer to the reference values than estimates from LG. These results are representative of those obtained for other branches (results not shown). The additional *χ*_1_ configuration information contained in RAM55 is thus allowing us to infer more-reliable phylogenies from alignments simulated under the 55-state model itself than does any of the 20-state models investigated.

**FIG. 6.**
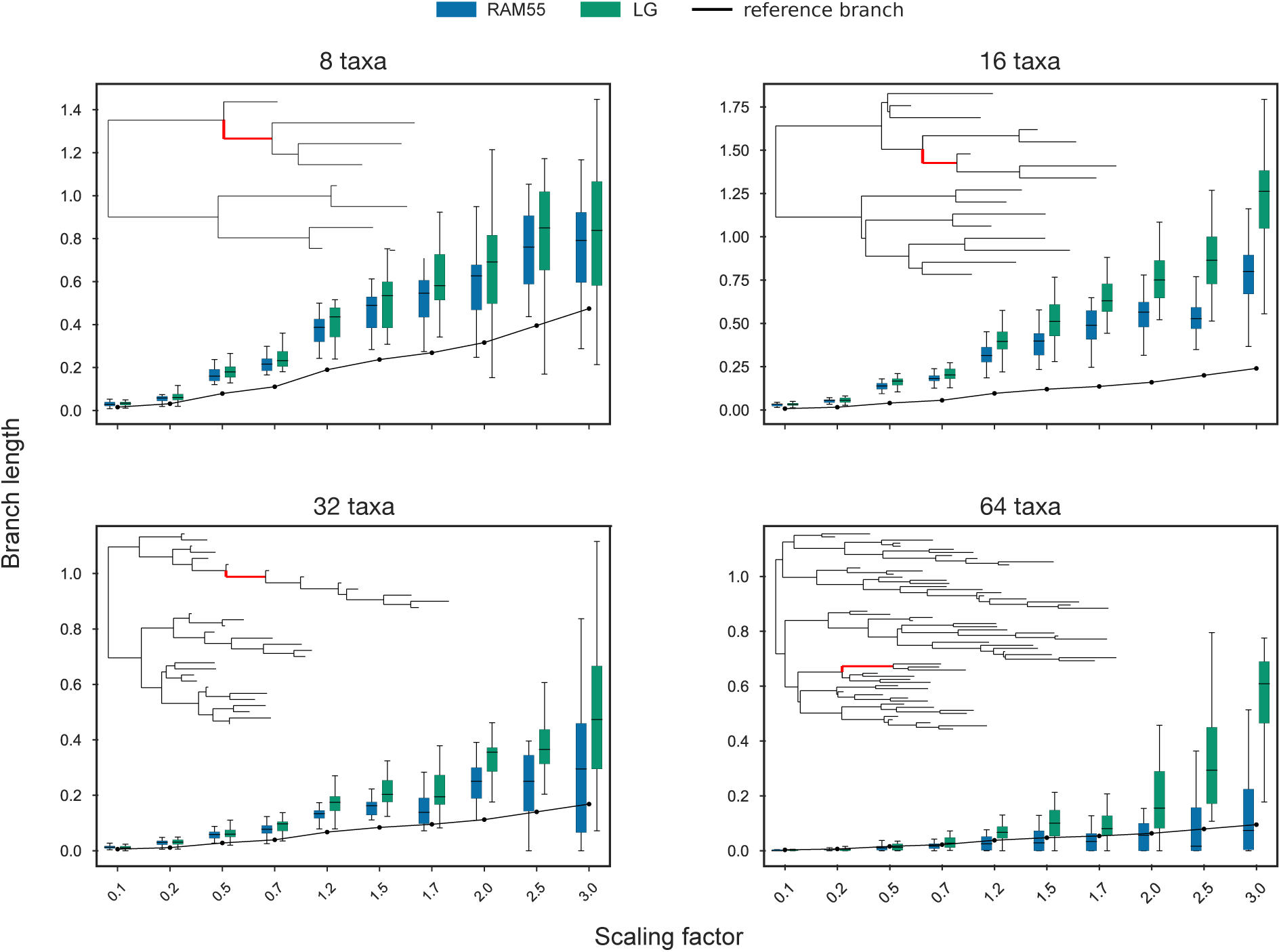
Individual branch length inference comparison. Trees inferred using RAM55 (blue) or LG (green), analyzing alignments (200 sites, 100 replicates per scaling) simulated using RAM55 and reference phylogenies of 8, 16, 32 and 64 taxa, scaled according to the factors displayed along the *x*-axis. Highlighted internal branches (indicated in red) have true lengths indicated by the solid lines; distributions of inferred lengths are shown as box-plots (evenly distributed horizontally and displaced for clarity).

### Model benchmarking: empirical alignments

We assessed RAM55’s performance on three empirical amino acid alignments — with 13, 82 and 46 taxa, respectively — for which we can obtain corresponding structural information (see *Empirical alignments*), and compared goodness-of-fit and inferred phylogenies across models. RAM55 was used in ML analyses, and results compared with those derived using structure-free models such as the 20-state models LG, WAG and our own RUM20, and the 55-state LGbyfreq-exp model which recognizes the frequencies of the 55 states but not their structural information content (see *Log-likelihood comparison across models, Likelihood calculation and maximization over phylogenies*).

Figure 7 shows the goodness-of-fit (measured by AIC values, see *Log-likelihood comparison across models*) for each empirical amino acid alignment under a variety of models. In all cases, RAM55 is a better fit for the data than all the other models used, indicated by the lower AIC values. Since our model is implemented in RAxML-NG (Kozlov *et al.*, 2018), it was also straightforward to incorporate a discrete gamma model of rate heterogeneity (Yang, 1993), maximum likelihood estimation of equilibrium frequencies from the observed data, or both in combination (see *Likelihood calculation and maximization over phylogenies*). The corresponding models, denoted RAM55+G, RAM55+F and RAM55+G+F, resulted in further improvements in the model’s fit, with RAM55+G+F performing best for all data sets. This empirical benchmark shows that RAM55 fits well when tested on three diverse datasets, and thus appears to be a valuable model of protein sequence evolution.

**FIG. 7.**
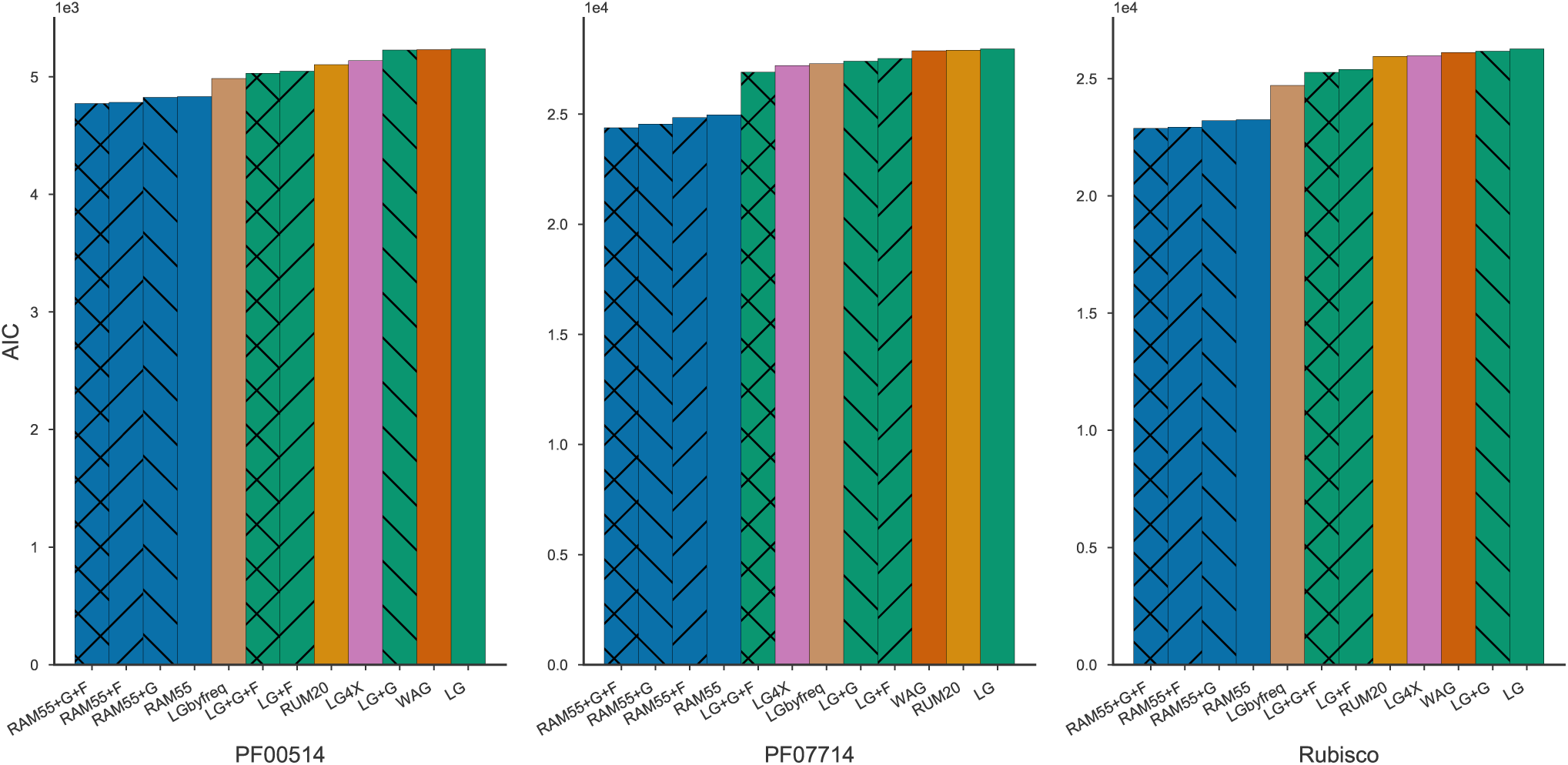
A comparison of RAM55 variants against other models in terms of Akaike information criterion (AIC) on three empirical rotasequence alignments: PF00514 (*β*-catenin-like repeat), PF07714 (tyrosine kinase) and rubisco. ‘+G’ models use a discrete gamma model of rate heterogeneity with 4 categories; ‘+F’ models use ML-estimated state frequencies estimated from the observed data.

### Ancestral state reconstruction

Having established RAM55’s ability to infer reliable phylogenies when structural information is available, we evaluate whether it can be used for two further tasks. The first is the reconstruction of ancestral amino acid states, which can also be achieved using a standard substitution model in which side chain configurations are not modelled. Second, we try to reconstruct the rotamer sequence in addition to the amino acid sequence, a capability which is unique to our model. We evaluated the performance on these tasks using both joint (Pupko *et al.*, 2000) and marginal (Yang *et al.*, 1995) reconstruction algorithms. The (55-state) RAM55 model can be applied to reconstruct plain (20-state) ancestral amino acid sequences, when present-day crystallography data are available, by first reconstructing ancestral rotasequences, and then simply masking the rotamer configuration information. The resulting ancestral amino acid sequences can then be compared to the known (masked) ancestor sequences in our simulations. We perform simulations as before under RAM55 using an 8-taxa reference topology, and then reconstruct ancestral amino acid states using RAM55 or LG and the joint reconstruction method (Pupko *et al.*, 2000). Terminal amino acid sequences (Fig. 8, nodes A and B) were also reconstructed in order to validate our “leave-leaves-out” (LLO) approach that serves as a proxy for ancestral sequence reconstruction when lacking a reference (see *Ancestral state reconstruction*). As shown in Fig. 8, it is possible to estimate terminal sequences with reasonable accuracy with this strategy (see also Sup. Figs. 6, 7), suggesting this is a viable method to evaluate reconstruction accuracy on empirical alignments, where ancestral reference sequences (and structures) are unlikely to be available. Figure 8 shows that our model performs equally or slightly better than LG in terms of of amino acid state reconstruction accuracy, particularly at longer evolutionary distances (see also Sup. Fig. 7, Sup. Tab. 2). Very similar results are achieved using the marginal reconstruction approach (Sup. Fig. 6, 8 and Sup. Tab. 2). These show that, in addition to exploiting information about *χ*_1_ configuration evolution to assist with model selection and phylogeny inference, RAM55 can be used to reconstruct ancestral sequences as well as a 20 state model, while as the same time providing information about the side-chain conformation which is not possible by any other method.

**FIG. 8.**
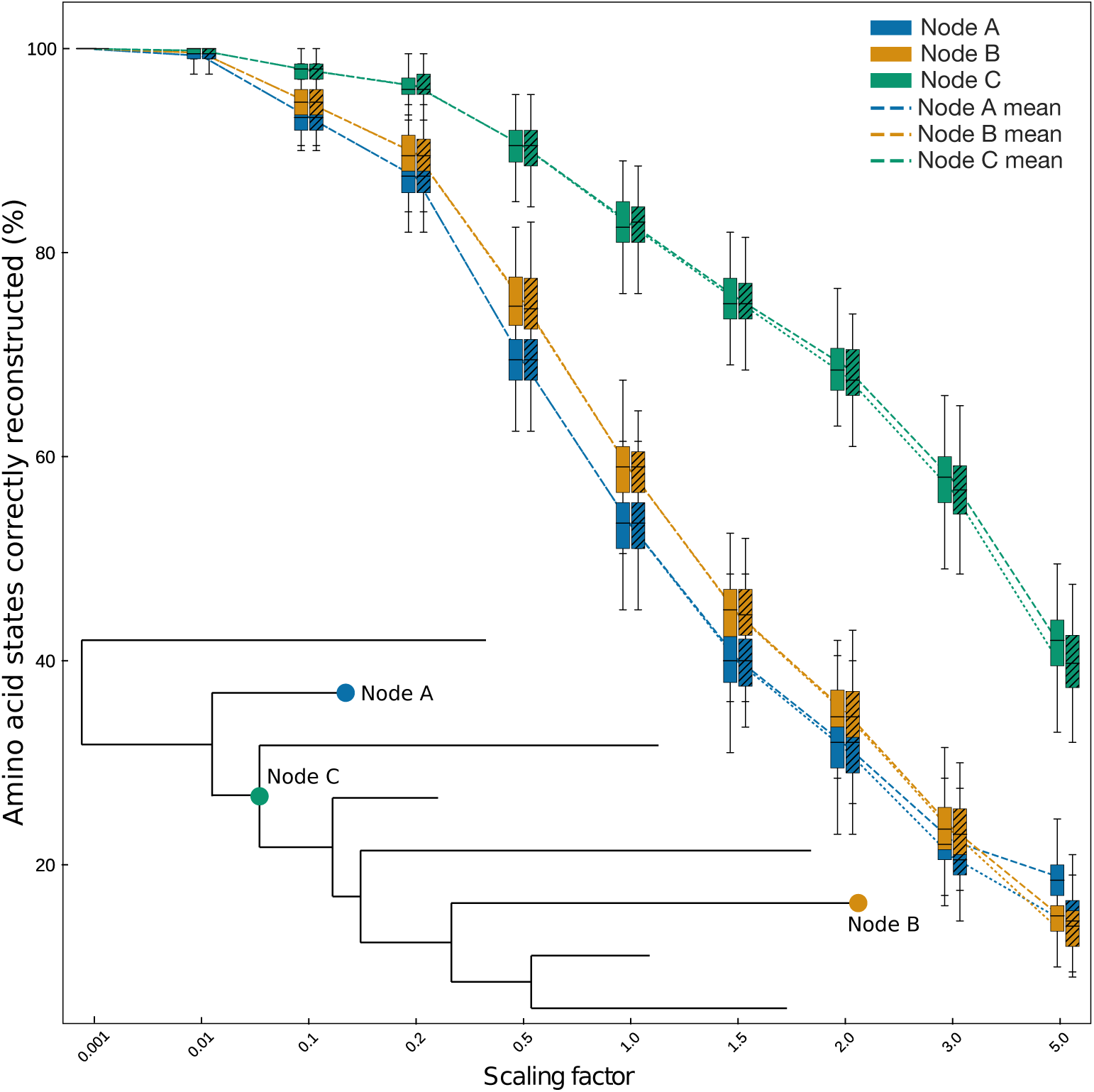
Amino acid reconstruction accuracy. Amino acid states inferred using joint reconstruction from rotasequence alignments (200 sites) simulated under RAM55 using our 8-taxon reference phylogeny and scaling its branches according to the factors reported on the *x*-axis (note the non-linear scale used for clarity). The joint reconstruction algorithm is then employed along with RAM55 and the true phylogeny to reconstruct rotasequences at various internal (C) or terminal (A, B) nodes. The *y*-axis indicates the proportion of amino acid states (i.e. masked rotasequence states) correctly reconstructed for each inferred sequence. Plain boxplots indicate the distribution of percentages of sites correctly reconstructed (*y*-axis) by this method; the same procedure is then repeated using LG on “masked” alignments (hatched boxplots). Each box-plot contains results from 100 simulation replicates for a given node.

We thus assessed our models’ accuracy when joint reconstructing ancestral rotamer states simulated under the model itself and our 8-taxa phylogeny. Figure 9 shows that RAM55 is able to infer the correct ancestral side chain configuration for residues belonging to internal sequences in almost all cases when the ancestral amino acid state is accurately reconstructed (as shown in Fig.8). Similar results are obtained for other alignment lengths and numbers of taxa in reference phylogenies (data not shown). These reconstructed ancestral rotamer states could be used to predict side-chain geometry for homology modelling of ancestral proteins, to assess which configuration better fits the evolutionary data.

**FIG. 9.**
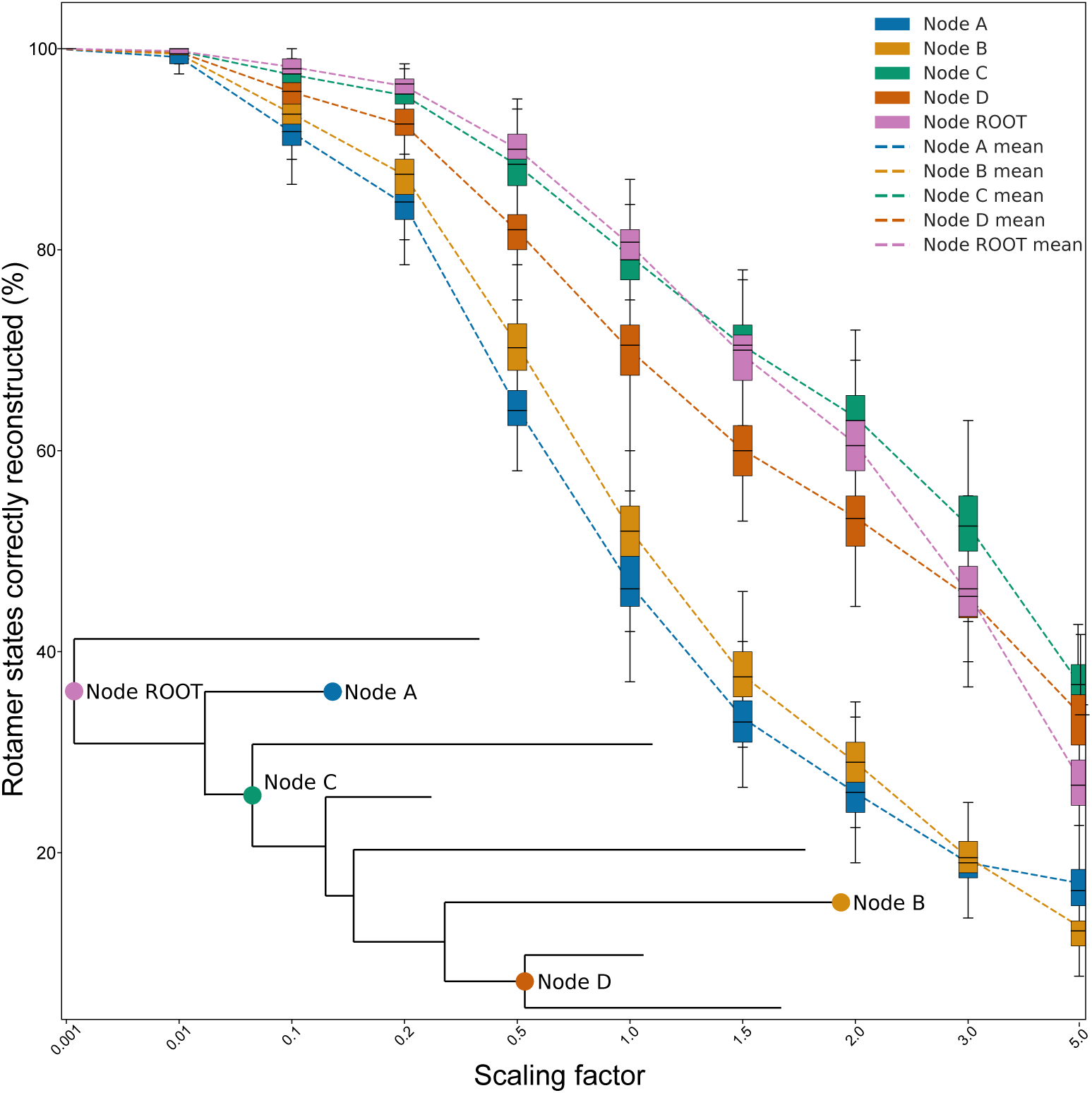
Rotamer state reconstruction accuracy. Rotamer states inferred using joint reconstruction from rotasequences (8 taxa, 200 sites) simulated under RAM55 using our 8-taxon reference phylogeny and scaling its branches according to the factors reported on the *x*-axis (note the non-linear scale used for clarity). The joint reconstruction algorithm is then employed along with RAM55 and the true phylogeny to reconstruct rotasequences at various internal (ROOT, C, D) or terminal (A, B) nodes. The *y*-axis indicates the proportion of rotamer states correctly reconstructed for each inferred sequence. Boxplots indicate the distribution of percentages of sites correctly reconstructed (*y*-axis) by this method. Each boxplot contains results from 100 simulation replicates for a given node.

## Conclusions

We have created a Dayhoff-like continuous-time Markov model that accounts for structural constraints on protein evolution by employing an expanded state set where each state corresponds to an amino acid along with its side-chain’s *χ*_1_ configuration. The exchange rates of our 55-state model, RAM55, clearly capture effects of local steric constraints, for example those dictating how an aromatic side-chain can be positioned without displacing or clashing with neighbouring residues. Other highly significant rotamer state exchange patterns, while still carrying valuable information for our model, have no obvious biochemical explanation and in some cases exhibit a tendency to avoid conserving *χ*_1_ configurations during amino acid exchanges. These exchange patterns deserve further exploration, perhaps by relating 3-D molecular descriptors to the exchange rates as has been attempted for amino acid exchange rates and 1-D biochemical properties (Grantham 1974, Dayhoff *et al.* 1978, Zoller and Schneider 2013).

Using simulated data, we confirmed that our 55-state model captures enough information to detect the *χ*_1_ configuration-aware expanded state space, and observed that it consistently offers detectably better fit to data compared to models that use the traditional 20-state space such as LG, WAG, and our RUM20. Further, RAM55 appears to infer equally or more reliable phylogenies than any of the 20-state models. This argues in favour of its consideration for phylogenetic analysis of protein sequences. Moreover, when applied to empirical data, the model provided a better fit than any of the traditional models evaluated. Our model can also be applied to perform structurally-aware reconstruction of ancestral sequences. Both amino acid and structural configuration states can be reliably inferred. Although there is little improvement in amino acid sequence reconstruction over traditional 20-state models, RAM55 could improve ancestral protein resurrection by (1) providing better phylogenies, which are valuable in themselves but also help towards (2) obtaining reconstructions of structural information, i.e. *χ*_1_ configurations, that are simply not possible by any other method. More generally, inferred rotamer states could be used to predict side-chain geometry for homology modelling, to assess which configuration better fits the evolutionary data. In this paper we apply our model to empirical data where both amino acid sequences and the corresponding high-quality atomic coordinates. While an increasingly large number of protein sequences can be associated with reliable X-ray crystallography data, and recent advances in cryo-electron microscopy promise to improve the resolutions that can be achieved for complex, dynamic molecular assemblies in their native state (Milne *et al.* 2013, Carroni and Saibil 2016, Venien-Bryan *et al.* 2017), many real-world applications of our approach might rely on data with a mixture of amino acid sequences and rotasequences, or amino acid sequences alone. Our model could be applied to this type of data by treating ambiguity regarding the rotameric state in the same manner in which sequencing errors and other forms of ambiguity are handled (Huelsenbeck 2002, Felsenstein 2004) thus allowing the information gained by accounting for the distinct evolutionary signatures of the rotamer states to be applied to all bioinformatics tasks relying on evolutionary modelling, as well as opening potential applications such as the prediction of side-chain geometries from amino acid sequences in the absence of other structural information.

## Materials and Methods

### Rotamer assignment and sequence alignments

In order to tabulate substitution events, a data set of aligned amino acid sequences annotated with their *χ*_1_ rotamer state was required. This was obtained from the Pfam database (Finn *et al.*, 2014) by first selecting those aligned sequences that are mapped to a high resolution crystal structure (<2.5Å) in the Protein Data Bank in Europe (Velankar *et al.*, 2010) to ensure we only retrieve those structures that are likely to be reliable. For each amino acid in each sequence, we assigned a *χ*_1_ rotamer configuration based on atomic coordinates, as defined in the Dunbrack rotamer library (Shapovalov and Dunbrack, 2011). We removed residues with an average B factor (Trueblood *et al.*, 1996) *>* 30 for the four atoms defining *χ*_1_ (*N, C*^*α*^, *C*^*β*^ and *C*^*γ*^ for most amino acids), to ensure that rotamer state assignments were based on unambiguous electron densities and not modelling artefacts. We also removed non-standard residues, disordered residues and those with peptide bonds exceeding 1.8Å, the last to ensure a continuous polypeptide. In this study we consider only *χ*_1_ configurations, and not those of rotatable bonds further along the side-chain, for a number of reasons: *χ*_1_ is present across all residues with the exception of glycine and alanine; it is closest to the backbone and thus usually better resolved in terms of atom positions; it conveys the most information about side-chain atom positions as all other side-chain atoms depend upon it; it gives us a manageable number of states; and it always connects two *sp*^3^ hybridised atoms, and thus is strictly rotameric and has exactly three conformational states (Dunbrack, 2002) although one is inaccessible in proline. These quality filtering steps resulted in alignments from 3,646 Pfam families, including 31,801 unique Uniprot entries, 251,194 PDBe structures and 81,523,991 residues.

### Tabulating substitution counts

We combined the amino acid sequences and rotamer state sequences to produce sequences in an expanded alphabet (see Table 1), which we refer to as ‘rotasequences’. Each rotasequence consists of symbol pairs (*A,R*), each of which specifies a state comprising the amino acid *A* (as employed by traditional 20-state models) and a *χ*_1_ rotamer configuration *R* (1, 2 or 3). For each family (see Supplementary Files) we then performed a sequence alignment-guided pairwise comparison of rotasequences. We used Pfam’s original domain alignment to construct a NJ phylogenetic tree (Saitou and Nei, 1987) using MAFFT (Katoh and Standley, 2013), and then iteratively tabulated differences between pairs of rotasequences by taking a circular tour through the NJ tree using an algorithm analogous to the one described by Korostensky and Gonnet (2000). While comparing pairs of rotasequences, we omitted those with a rotasequence identity < 75%, to minimise the risk of multiple substitution events at the same site being tabulated as a single observed difference. This approach results in an efficient set of pairwise comparisons using all leaves of the trees with each observed difference counted at most twice. We tabulated 30,439,912 counts, corresponding to 4,508,390 rotamer state substitutions and 25,931,520 instances of rotamer state conservation.

We then computed the observed number of occurrences of sites in all aligned sequence pairs with rotamer states (*A,R*) in one sequence and (*A*′*,R*′) in the other as *n*_(*A,R*),(*A*′*,R*′)_. While these counts could be used directly to calculate an instantaneous rate matrix (IRM), this would result in biases arising from the filtering procedures described above. For example, because alanine and glycine can never be filtered out by B factor, these residues are overrepresented. Further, some amino acids, such as those commonly well-packed into the core of the protein, are better resolved and have lower B factors than those more commonly found at the protein surface, and are thus also overrepresented (Sup. Fig. 9). To account for this we also compute substitution event counts (*n*_*A, A*′_) for a Dayhoff-like 20-state empirical model (RUM20) using the same Pfam-based dataset, but ignoring *χ*_1_ configurations and without performing B factor filtering. Our normalized rotamer state exchange count matrix 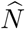, recovering the actual observed residue frequencies, then becomes:

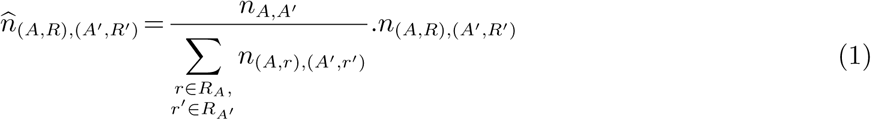

where *R*_*A*_ = {*R* : (*A,R*) ∈ *S*_55_} is the set of rotamer configurations *R* such that the corresponding pair (*A,R*) is a member of *S*_55_, the set of all 55 possible combinations shown in Table 1.

The 55-state 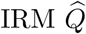 is computed from these normalized counts as described by Kosiol and Goldman (2005): the instantaneous rate of change of (*A,R*) into (*A* ′*,R* ′) (with (*A,R*) ≠(*A* ′*,R* ′)) is given by the number of such events as a proportion of all observations of (*A,R*):

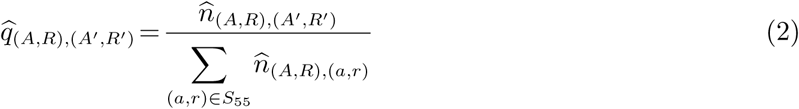

As usual, diagonal elements 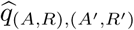 are set so that row sums of the IRM equal 0 (Kosiol and Goldman, 2005). The 20-state RUM20 IRM was similarly obtained from the unfiltered 20-state counts *n*_*A,A*′_, for comparative purposes.

### Rate scaling

Times and branch lengths are typically measured as the expected number of substitutions per site (Felsenstein, 2004). Our rates were therefore first scaled according to *ρ* so that, at equilibrium, they will result on average in one rotamer state ((*A,R*) → (*A*′*,R*′) with {(*A,R*),(*A*′*,R*′)} ∈ *S*_55_) substitution per unit of time (Lio and Goldman, 1998):

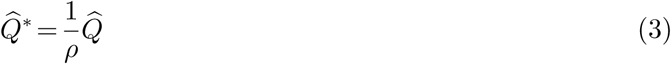

With

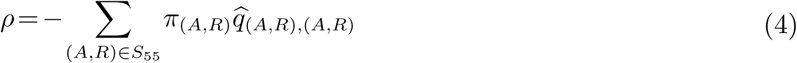

where *π*_(*A,R*)_ is the equilibrium frequency of rotamer state (*A,R*) obtained from the normalized counts 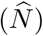. Because we have an expanded state set, 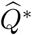 results in one rotamer state substitution per unit time but less than one amino acid state substitution. We therefore perform a further scaling step in order to allow direct comparison of branch lengths estimated with our 55-state model and any 20-state model, i.e. in terms of number of amino acid state substitutions. This additional scaling factor *ρ*\* is defined as:

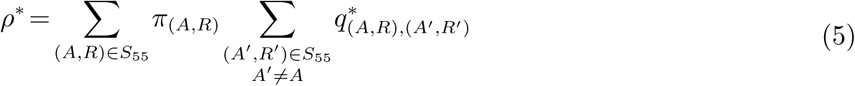

and corresponds to the proportion of rotamer state changes where the amino acid changes, irrespective of *χ*_1_ configuration. Then the ‘superscaled’ IRM is given by:

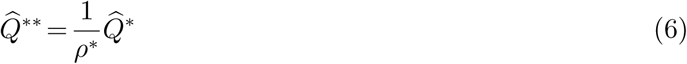

This matrix, at equilibrium, has on average 1*/ρ*\* = 1.79 state changes per unit time, consisting of 1 amino acid state substitution plus 0.79 (i.e. (1-*ρ*\*)*/ρ*\*) *χ*_1_ configuration changes that are invisible to traditional 20×20 models. This means that branch lengths are directly comparable to those under 20-state structure-free models that can only detect amino acid changes. RAxML-NG’s implementation of RAM55 provides output in these superscaled time units (comparable to traditional amino acid distances) and also in the units of eqn. (3).

From 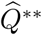, final scaled exchangeabilities (available in Sup. Files) were obtained as:

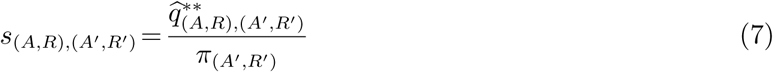

(Liò *et al.* 1998, Whelan and Goldman 2001). Exchangeabilities computed from a general dataset can be combined with state frequencies estimated from any particular dataset under study, and a dataset-specific IRM can thus be obtained by inverting eqn. (7). This hybrid parametrization procedure, denoted by adding ‘+F’ to a model name, can produce a significant improvement in model fit (Thorne and Goldman 2007, Perron *et al.* ressin press).

### Rotamer state exchangeability analysis

Each pair of different amino acids corresponds to a 3×3 sub-matrix (with the exception of pairs including alanine, glycine and proline) in 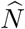. Since 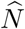 is symmetric there are 136 unique 3×3 sub-matrices. For each of these we computed the Pearson’s *χ*2 statistic and *p*-value (with Bonferroni correction) for the hypothesis test of independence of the observed *χ*_1_ rotamer configuration change frequencies, where the expected frequencies are computed based on the marginal sums under the assumption of independence. Pairs of residues with Bonferroni *p*-value < 0.05 show significant association among their rotamer states; only these are considered for further analysis (e.g. Fig. 3).

We assessed the strength of association between the *χ*_1_ rotamer configurations of each pair of residues using Cramér’s 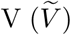 with bias correction (Bergsma, 2013). We also computed the proportion of counts that lie on each 3×3 sub-matrix’s diagonal (‘diagonal ratio’). The latter is a measure of the tendency of a pair of exchanging residues to conserve their *χ*_1_ rotamer configuration. To better assess trends in diagonal ratio and 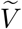, residues with three available *χ*_1_ rotamer configurations (excluding methionine) are then classified into six groups depending on the biochemical properties of their side-chains: aliphatic (isoleucine, leucine, valine), aromatic (phenylalanine, tryptophan, tyrosine), positive (arginine, lysine, histidine), carboxylamine (asparagine, glutamine), negative (aspartic acid, glutamic acid) and hydroxyl (serine, threonine). While this is an imperfect classification, as residues do not fit unambiguously into distinct, discrete groups and have multiple salient features, it nonetheless helps up to better visualize how side-chain properties influence exchange rates.

### Overlap of backbone distributions

The global structure of a protein is determined largely by the configuration of the peptide backbone onto which the side-chains are bonded, which can be characterised by the dihedral angle of the two rotatable bonds, *ϕ* and *ψ*, of each amino acid. Steric effects determine which combinations of *ϕ* and *ψ* are allowed, and which are favoured, as commonly visualised using a Ramachandran plot, which shows permitted regions and observed distributions over *ϕ* and *ψ* (Ramachandran *et al.*, 1963). As these steric effects arise, in part, from the side-chain, and the rotameric states of the side chain are influenced by the conformation of the backbone, each rotamer state has its own probability distribution on the Ramachandran plot (Dunbrack and Karplus, 1993). To test the hypothesis that rotamer states preferentially exchange with other rotamer states with similar backbone dependencies, we calculated the overlap between the (*ϕ,ψ*) distributions of pairs of rotamer states (*i, j* ∈ *S*_55_) from their probability density functions, *f* • (*ϕ,ψ*), as estimated from rotamer counts in the Dunbrack backbone-dependant rotamer library (Shapovalov and Dunbrack, 2011), using the equation

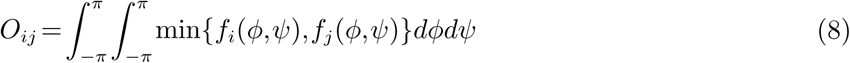

### Kullback-Leibler divergence

We measured the amount of information lost, regarding the amino acid sequence, when a 20-state model is used to approximate our 55-state model by computing the Kullback-Leibler (KL) divergence in *bits* (Kullback and Leibler, 1951) for each rotamer state (*A,R*) and its corresponding amino acid state *A*. This metric measures the divergence between the amino acid probability distribution at time *t*, when starting with rotamer state (*A,R*), between the RAM55 model in which both *A* and *R* and considered and the RUM20 model in which only *A* is used. The KL divergence is computed as a function of evolutionary time *t* using:

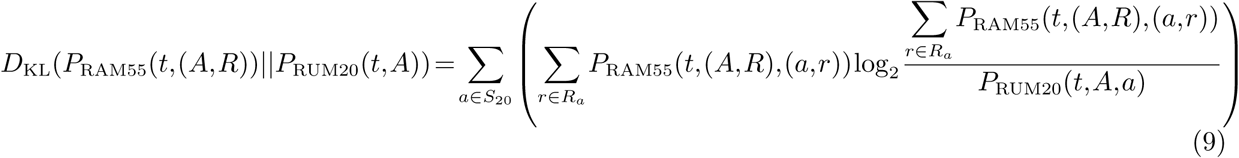

with *S*_55_ and *S*_20_ being the 55- and 20-state spaces, *R*_*a*_ the *χ*_1_ configurations of amino acid *a, P*_RUM20_(*t*) and *P*_RAM55_(*t*) the probability matrices of the respective models at time *t* (see eqn. (10) below), and (e.g.) *P*_RAM55_(*t*,(*A,R*),(*a,r*)) the ((*A,R*),(*a,r*)) element of *P*_RAM55_(*t*).

### Likelihood calculation and maximization over phylogenies

We implement our models using maximum likelihood (ML) methods applied to multiple sequence alignments (Felsenstein, 2004). This standard approach searches for the tree *T* that maximizes the likelihood function with substitutions being modelled by a Markov process. Markovian state substitutions over time *t* are described by a probability matrix defined by

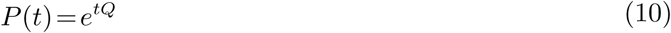

where *Q* is the IRM of the Markov process. The likelihood of *T* (including tree topology and branch lengths) given data (alignment) *X* and IRM *Q* can then be computed as:

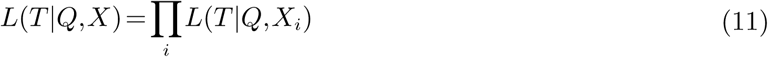

where *L*(*T* |*Q,X*_*i*_) corresponds to the likelihood of *T* given the states observed at site *i* of *X* (site independence assumption). *L*(*T* |*Q,X*_*i*_) is computed by applying eqn. (10) to each tree branch and using the pruning algorithm (Felsenstein, 1981). Maximizing *L* over *T* provides estimates 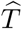 and thus the most-likely phylogeny given the observed data and the current substitution model. The ‘+F’ approach to matching the model’s state frequencies to the observed data can be implemented by simultaneously maximizing *L* over these frequencies.

It is also generally acknowledged that sites do not evolve at the same rate, due to various evolutionary constraints. The most common way of accounting for this heterogeneity is to assume that rates across sites follow a discretized gamma distribution (Yang, 1994). The shape parameter of the gamma distribution, *α*, is usually estimated by ML along with *T* as it is considered specific to each dataset. Models using the gamma distribution to model rate heterogeneity are denoted ‘+G’.

In this study, all ML tree inferences were performed using RAxML-NG (Kozlov *et al.*, 2018) which, following the needs of this study, now has functionality allowing custom state spaces and rate matrices of any size, permitting us to use our 55-state model RAM55 to infer tree topology, branch lengths and likelihoods that can be used for model fitting and comparisons. It also permits the +F and +G variants of substitution models through its “+FO” and “+G” options. Our expanded state-space has some inevitable repercussions for CPU time, not least because 20-state models benefit from a highly-optimized likelihood computation in RAxML-NG, whereas the RAM55 model currently works with general kernels that are less efficient. Nevertheless, computation times remain acceptable, tending to be 5–10 times longer than using 20-state models (Sup. Fig. 10).

### Tree generation and alignment simulation

We simulated sequence alignments under RAM55 using four randomly generated trees (8, 16, 32 or 64 taxa; branch lengths ∈ [0.01,0.5]; see Sup. Fig. 11 and Sup. Files) as guide and a substitution simulation approach based on Method 1 of Fletcher and Yang (2009), modified for our expanded state-set. Additionally, a pruned (mammals) and scaled version of the Ensembl-compara species tree (Herrero *et al.* 2016, see Sup. Files) was also used. To allow investigation of a realistic range of sequence divergences (around 10–85%) while maintaining consistent tree topologies, all branches of our trees were scaled according to a set of 10 scaling factors: {0.1, 0.2, 0.5, 0.7, 1.2, 1.5, 1.7, 2, 2.5, 3} for model benchmarking simulations, or {0.001, 0.01, 0.1, 0.2, 0.5, 1, 1.5, 2, 3, 5} for ancestral reconstruction simulations. For each scaled tree we generated 100 rotasequence alignments of a realistic length (200 or 1,000 sites) using the strategy detailed above. These combinations of simulation parameters were designed to generate a broad range of evolutionary scenarios that might be encountered in empirical studies.

Under 55-state models, simulated alignments were analysed in this form; their constituent rotasequences were converted into amino acid sequences for inference under 20-state models by ‘masking’ the rotamer configuration component of their rotamer states (i.e. (*A,R*) → *A*).

### Log-likelihood comparison across models

When selecting the best fitting model for a specific dataset, an information-theoretic score such as the Akaike Information Criterion (AIC) is frequently used (Akaike 1974, Sullivan and Joyce 2005). This approach fits well with our comparison where many models of interest are non-nested and, in cases, have different state spaces. The AIC score is defined as

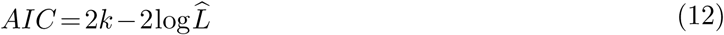

where *k* is the number of estimated parameters in the model and 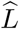 is the maximized value of the likelihood function of eqn. (11). However, the likelihood function depends upon a model’s state-space (Anderson and Burnham, 2002): 20-state models of amino acid substitution cannot be directly compared with our 55-state model as they exist in different state-spaces. Whelan and colleagues have developed a generalized ‘correction’ allowing the comparison of likelihoods between state-spaces (Whelan *et al.*, 2015). This strategy is applicable to any two state-spaces (*D,C*) providing that (1) each state in *D* maps to a single state in *C* and each state in *C* maps to a unique set of states in *D*, and (2) both likelihoods are obtained from the same original alignments, *X*^*C*^ and *X*^*D*^, with *X*^*C*^ being the “compounded” version of *X*^*D*^ following the state mapping. The corrected likelihood of the distinct model (*D*) can then be expressed in terms of the compound model (*C*) likelihood and an adapter function as:

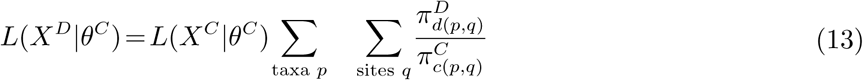

where *θ*^*C*^ and *θ*^*D*^ represent the totality of parameters from *C* and *D*; *d*(*p,q*) and *c*(*p,q*) are the distinct and compound states observed for taxon *p* at site *q*; and 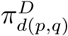 and 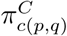 are these states’ equilibrium frequencies in their respective substitution models. In our application of this approach the distinct model *D* corresponds to RAM55, whose states can be uniquely compounded into amino acid states (e.g. TRP3 → TRP), and the compound model *C* corresponds to a 20-state amino acid model (e.g. WAG, LG or our RUM20) whose states can be mapped to a unique set of rotamer states (e.g. TRP → {TRP1, TRP2, TRP3}).

As an independent approach to test the contribution of knowledge of rotameric configuration-state substitutions, we generated 55-state models that were expanded versions of the 20-state LG model (Le and Gascuel, 2008). Likelihoods are directly comparable to RAM55’s since they share the same state-space. This model expansion operation was performed with the introduction of no information about the additional states (LGexp model) or, alternatively, by accounting for just the observed frequencies of these additional states in our dataset (LGbyfreq-exp). In each case, we started from LG’s exchangeabilities and reconstructed a raw substitution count matrix by reversing 20-state versions of eqns. (7) and (2). For LGexp, this reconstructed counts matrix *N* was then expanded into a 55-state counts matrix 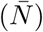 according to

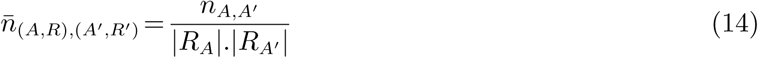

where |*R*_*A*_|.|*R*_*A*′_ | is the product of the dimensions of a sub-matrix in 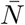 corresponding to a single cell of *N*. Eqns. (2) and (7) are then applied to 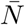 to derive the IRM for the LGexp model. This expanded model represents the ‘most-uninformed’ expression of a 20-state model in a 55-state space, introducing rotamer states but no information about their relative frequencies or replacement rates.

Alternatively, for LGbyfreq-exp, *N* was expanded according to

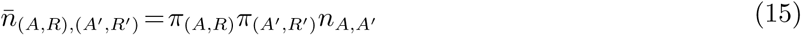

where *π*_(*A,R*)_*π*_(*A*′*,R*′)_ is the product of RAM55’s equilibrium frequencies for states (*A,R*) and (*A*′*,R*′). The LGbyfreq-exp expanded model’s rates are therefore informed about each rotamer state’s frequency, but not the relative rates of replacement between them observed in real protein sequences. We can thus compare all our models in term of their fit for a specific dataset using eqn. (12) with the likelihood term corresponding either simply to 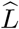 for 55-state models (RAM55, LGexp, LGbyfreq-exp) or to the state-corrected likelihood obtained from eqn. (13) for 20-state models (RUM20, LG, WAG). The latter is referred to as a ‘state-corrected AIC score’.

### Empirical alignments

We assessed RAM55’s goodness-of-fit and performance on empirical data using rotasequence alignments (available in Sup. Files) that can be masked by removing the rotamer configuration information in order to convert then to amino acid sequences for comparison inferences with 20-state models. Alignments PF00514 and PF07714 correspond to two Pfam family alignments and their corresponding structural information from PDBe: *β*-catenin-like repeat and tyrosine kinase, respectively. We followed the same procedure previously used (see *Rotamer assignment and sequence alignments)* to assign rotamer states, using Pfam’s domain alignment and mapping of sites to PDBe residues. These alignments are relatively short — 13 taxa and 334 sites for PF0054, 82 taxa and 345 sites for PF07714 — as they only include those portions of sequences Pfam recognizes as part of that family’s domain. The third aligment was obtained by querying Uniprot (UniProt Consortium, 2017) with the term “rubisco” and obtaining the corresponding PDBe entries with no reliance on Pfam domain alignments. Rotamer states were then assigned as described when estimating RAM55; however, in this case we did not limit ourselves to Pfam’s definition of a family domain and this results in a longer alignment (46 taxa, 681 sites).

### Ancestral state reconstruction

There are two broad categories of approaches to the problem of ancestral sequence reconstruction. Marginal reconstruction assigns the most likely state to each ancestral sequence at a given site independently of the states reconstructed for other ancestral sequences at that site. Joint reconstruction instead finds an assignment of ancestral states throughout the tree that jointly maximizes the likelihood of the observed data at that site (Yang *et al.* 1995, Yang 2007). We used both the marginal reconstruction algorithm (Yang *et al.*, 1995) and Pupko *et al.*’s implementation of the joint reconstruction algorithm (Pupko *et al.*, 2000) to infer ancestral rotamer and amino acid sequences; we adapted both algorithms to fit our expanded state-space.

To test whether our RAM55 model allowed us to correctly infer unobserved ancestral states starting from data simulated under the model itself, rotasequence alignments was simulated using RAM55, trees with fixed topology (Sup. Fig. 11) and branch lengths scaled, in turn, according to a set of factors: {0.001, 0.01, 0.1, 0.2, 0.5, 1, 1.5, 2, 3, 5}. For each scaled tree, 100 replicate alignments were generated: these included internal node rotasequences to be used as references against which inference accuracy was assessed. Masked versions of all sequences were also created, to allow 20-state model inference (i.e. inference of ancestral amino acid sequence alone) using LG. The phylogenies from these simulations were then employed alongside RAM55 or LG to reconstruct ancestral states. Finally, reconstructed rotasequences or amino acid sequences were compared position-by-position against the simulated reference sequences and the results reported in terms of percent sequence identity (percent correct inference).

We then investigated RAM55’s performance when reconstructing ancestral rotasequences or amino acid sequences from empirical rotasequences. Ideally this would be performed by comparison of inferred ancestral rotasequences with known ancestral structures. While an increasing number of resurrected ancestral protein structures have been resolved (e.g. Konno *et al.* 2011, Ingles-Prieto *et al.* 2013, Hart *et al.* 2014, Risso *et al.* 2014, Clifton *et al.* 2018), their rarity, combined with the fact that most of these studies reconstruct ancestral amino acid sequences from alignment of present-day proteins that in many cases lack high quality structural information, do not allow a systematic comparison of our reconstructed rotasequences with reference ancestral rotasequences obtained from deposited structures. To overcome this, we employed a “leave-leaves-out” (LLO) approach (Sup. Fig. 12) in which we remove a pair of terminal sibling nodes from the alignment and proceed to reconstruct all internal nodes including one of the aforementioned pair of taxa according to the marginal or joint algorithms. (Pairs of sibling terminal nodes (A, B), as opposed to single terminal nodes (A), were removed as otherwise a remaining close neighbour of A could allow for easy reconstruction of A’s sequence.) LLO allows us to compare the inferred terminal sequence against the known original, as a proxy for the desired comparison. This approach was first validated on terminal sequences simulated under RAM55 and then used on empirical sequences from the PF00514 alignment; in this case the phylogeny inferred using RAxML-NG and RAM55 is used for the reconstruction process along with RAM55 or LG.

### Code availability

Code used to generate random trees and simulate substitutions along their branches (see *Tree generation and alignment simulation*) is available at: https://bitbucket.org/uperron/ram55. This repository also includes our implementations of the joint and marginal reconstruction algorithms (see *Ancestral state reconstruction*), as modified for our expanded state-set. In addition, we provide an example of how to obtain a rotasequence alignment, suitable for tree inference with RAxML-NG and RAM55, using a user-submitted set of Uniprot IDs and the PDBe API.

## Acknowledgments

We thank Nicola De Maio and Claudia Weber at EMBL-EBI who provided insight and expertise that greatly assisted this work. This work was supported by EMBL (U.P., N.G.), the Klaus Tschira foundation (A.M.K., A.S.) and the BBSRC (Future Leader Fellowship BB/N011600/1, to I.H.M.).

## Supplementary Tables

**Supplementary Table 1:** Best model (AIC) for each category of simulated alignment. 1000-site alignments are simulated under the RAM55 model and various randomly-generated reference phylogenies (4,8,16,32 and 64 taxa) scaled according to a set of scaling factors. For each phylogeny and scaling factor pair the table reports the proportion of 100 replicates where each model achieves the lowest AIC (or state-corrected AIC, see *Log-likelihood comparison across models*) when compared against all other models.

| Taxa | Scaling factor | RAM55 | RUM20 | LG | LGexp | LGbyfreq-exp | LG+F |
| --- | --- | --- | --- | --- | --- | --- | --- |
| 4 | 0.1 | 1.00 | 0.00 | 0.0 | 0.0 | 0.0 | 0.0 |
| 4 | 0.2 | 1.00 | 0.00 | 0.0 | 0.0 | 0.0 | 0.0 |
| 4 | 0.5 | 1.00 | 0.00 | 0.0 | 0.0 | 0.0 | 0.0 |
| 4 | 0.7 | 1.00 | 0.00 | 0.0 | 0.0 | 0.0 | 0.0 |
| 4 | 1.2 | 1.00 | 0.00 | 0.0 | 0.0 | 0.0 | 0.0 |
| 4 | 1.5 | 0.96 | 0.04 | 0.0 | 0.0 | 0.0 | 0.0 |
| 4 | 1.7 | 0.99 | 0.01 | 0.0 | 0.0 | 0.0 | 0.0 |
| 4 | 2.0 | 0.96 | 0.04 | 0.0 | 0.0 | 0.0 | 0.0 |
| 4 | 2.5 | 0.93 | 0.07 | 0.0 | 0.0 | 0.0 | 0.0 |
| 4 | 3.0 | 0.77 | 0.23 | 0.0 | 0.0 | 0.0 | 0.0 |
| 8 | 0.1 | 1.00 | 0.00 | 0.0 | 0.0 | 0.0 | 0.0 |
| 8 | 0.2 | 1.00 | 0.00 | 0.0 | 0.0 | 0.0 | 0.0 |
| 8 | 0.5 | 1.00 | 0.00 | 0.0 | 0.0 | 0.0 | 0.0 |
| 8 | 0.7 | 1.00 | 0.00 | 0.0 | 0.0 | 0.0 | 0.0 |
| 8 | 1.2 | 0.99 | 0.01 | 0.0 | 0.0 | 0.0 | 0.0 |
| 8 | 1.5 | 0.99 | 0.01 | 0.0 | 0.0 | 0.0 | 0.0 |
| 8 | 1.7 | 0.99 | 0.01 | 0.0 | 0.0 | 0.0 | 0.0 |
| 8 | 2.0 | 0.96 | 0.04 | 0.0 | 0.0 | 0.0 | 0.0 |
| 8 | 2.5 | 0.86 | 0.14 | 0.0 | 0.0 | 0.0 | 0.0 |
| 8 | 3.0 | 0.89 | 0.11 | 0.0 | 0.0 | 0.0 | 0.0 |
| 16 | 0.1 | 1.00 | 0.00 | 0.0 | 0.0 | 0.0 | 0.0 |
| 16 | 0.2 | 1.00 | 0.00 | 0.0 | 0.0 | 0.0 | 0.0 |
| 16 | 0.5 | 1.00 | 0.00 | 0.0 | 0.0 | 0.0 | 0.0 |
| 16 | 0.7 | 1.00 | 0.00 | 0.0 | 0.0 | 0.0 | 0.0 |
| 16 | 1.2 | 1.00 | 0.00 | 0.0 | 0.0 | 0.0 | 0.0 |
| 16 | 1.5 | 1.00 | 0.00 | 0.0 | 0.0 | 0.0 | 0.0 |
| 16 | 1.7 | 0.95 | 0.05 | 0.0 | 0.0 | 0.0 | 0.0 |
| 16 | 2.0 | 0.95 | 0.05 | 0.0 | 0.0 | 0.0 | 0.0 |
| 16 | 2.5 | 0.84 | 0.16 | 0.0 | 0.0 | 0.0 | 0.0 |
| 16 | 3.0 | 0.80 | 0.20 | 0.0 | 0.0 | 0.0 | 0.0 |
| 32 | 0.1 | 1.00 | 0.00 | 0.0 | 0.0 | 0.0 | 0.0 |
| 32 | 0.2 | 1.00 | 0.00 | 0.0 | 0.0 | 0.0 | 0.0 |
| 32 | 0.5 | 1.00 | 0.00 | 0.0 | 0.0 | 0.0 | 0.0 |
| 32 | 0.7 | 1.00 | 0.00 | 0.0 | 0.0 | 0.0 | 0.0 |
| 32 | 1.2 | 1.00 | 0.00 | 0.0 | 0.0 | 0.0 | 0.0 |
| 32 | 1.5 | 1.00 | 0.00 | 0.0 | 0.0 | 0.0 | 0.0 |
| 32 | 1.7 | 1.00 | 0.00 | 0.0 | 0.0 | 0.0 | 0.0 |
| 32 | 2.0 | 1.00 | 0.00 | 0.0 | 0.0 | 0.0 | 0.0 |
| 32 | 2.5 | 1.00 | 0.00 | 0.0 | 0.0 | 0.0 | 0.0 |
| 32 | 3.0 | 0.98 | 0.02 | 0.0 | 0.0 | 0.0 | 0.0 |
| 64 | 0.1 | 1.00 | 0.00 | 0.0 | 0.0 | 0.0 | 0.0 |
| 64 | 0.2 | 1.00 | 0.00 | 0.0 | 0.0 | 0.0 | 0.0 |
| 64 | 0.5 | 1.00 | 0.00 | 0.0 | 0.0 | 0.0 | 0.0 |
| 64 | 0.7 | 1.00 | 0.00 | 0.0 | 0.0 | 0.0 | 0.0 |
| 64 | 1.2 | 1.00 | 0.00 | 0.0 | 0.0 | 0.0 | 0.0 |
| 64 | 1.5 | 1.00 | 0.00 | 0.0 | 0.0 | 0.0 | 0.0 |
| 64 | 1.7 | 1.00 | 0.00 | 0.0 | 0.0 | 0.0 | 0.0 |
| 64 | 2.0 | 1.00 | 0.00 | 0.0 | 0.0 | 0.0 | 0.0 |
| 64 | 2.5 | 1.00 | 0.00 | 0.0 | 0.0 | 0.0 | 0.0 |
| 64 | 3.0 | 1.00 | 0.00 | 0.0 | 0.0 | 0.0 | 0.0 |

**Supplementary Table 2:** Empirical amino acid reconstruction accuracy. Amino acid state reconstruction from the PF00514/*β*-catenin-like repeat alignment using RAM55 or LG and either the joint or marginal reconstruction algorithms. Scores represent the percentage of sites correctly reconstructed. Reconstruction was limited to terminal nodes using the LLO approach (see *Ancestral state reconstruction*), due to lack of a reference for internal sequences.

| Taxa | RAM55 joint | LG joint | RAM55 marginal | LG marginal |
| --- | --- | --- | --- | --- |
| Q71VM4/147-186 | 75.15% | 76.05% | 72.75% | 72.16% |
| P52292/410-447 | 76.35% | 75.75% | 73.05% | 73.05% |
| P52294/158-197 | 80.84% | 80.24% | 76.95% | 76.35% |
| P52293/251-279 | 81.14% | 80.84% | 78.44% | 77.55% |
| Q9C2K9/158-194 | 89.22% | 89.82% | 81.74% | 81.44% |
| O60684/113-154 | 88.92% | 88.32% | 85.03% | 85.03% |
| P35222/229-262 | 89.22% | 89.22% | 89.52% | 89.22% |
| P25054/649-689 | 88.92% | 88.32% | 89.22% | 89.52% |
| Q02821/372-411 | 97.01% | 97.01% | 97.31% | 96.71% |
| Q02248/229-262 | 96.41% | 96.41% | 97.01% | 96.41% |
| Q99959/386-423 | 95.51% | 95.51% | 95.81% | 95.81% |
| O60716/440-481 | 96.41% | 96.41% | 97.31% | 97.31% |
| O44326/361-402 | 99.40% | 99.10% | 99.40% | 99.40% |

## Supplementary Figures

**Supplementary Figure 1:**
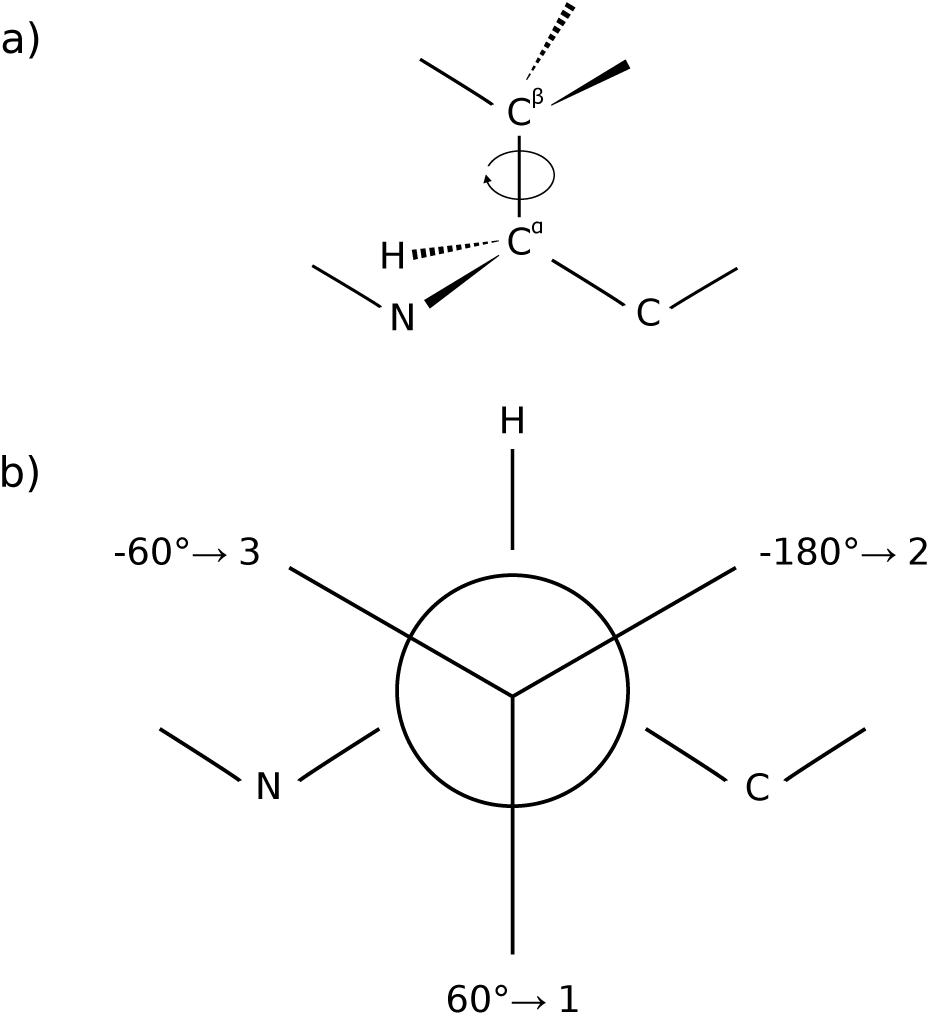
Definition of rotamer configurations. **(a)** A *χ*_1_ rotamer configuration is defined by the dihedral angle generated by the rotation of the *C*^*α*^ - *C*^*β*^ bond (curved arrow). **(b)** The three stable configurations correspond to specific *χ*_1_ dihedral angle values: ∼60°for configuration 1, ∼-180°for configuration 2 and ∼-60°for configuration 3. (See also Fig. 1 and Table 1)

**Supplementary Figure 2:**
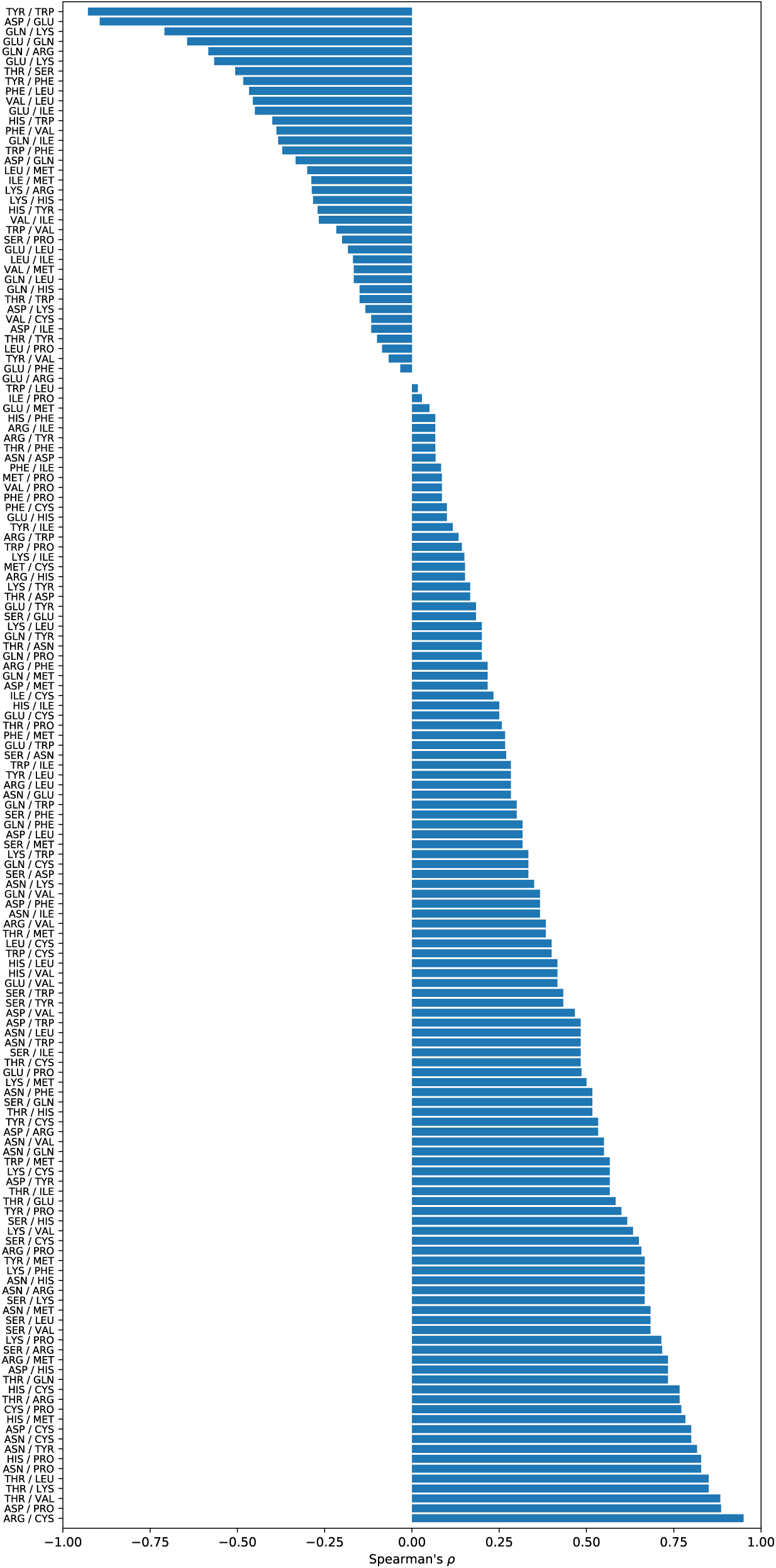
Correlation between exchangeability and the overlap between (*ϕ,ψ*) distributions. For each amino acid pair (excluding alanine and glycine), the correlation between the exchangeabilities of their *χ*_1_ configurations and the overlap between their Ramachandran probability distributions is shown. The preponderance of positive correlations indicates that, for most amino acid pairs, there is a tendency to exchange between side-chain geometries that accommodate similar backbone geometries.

**Supplementary Figure 3:**
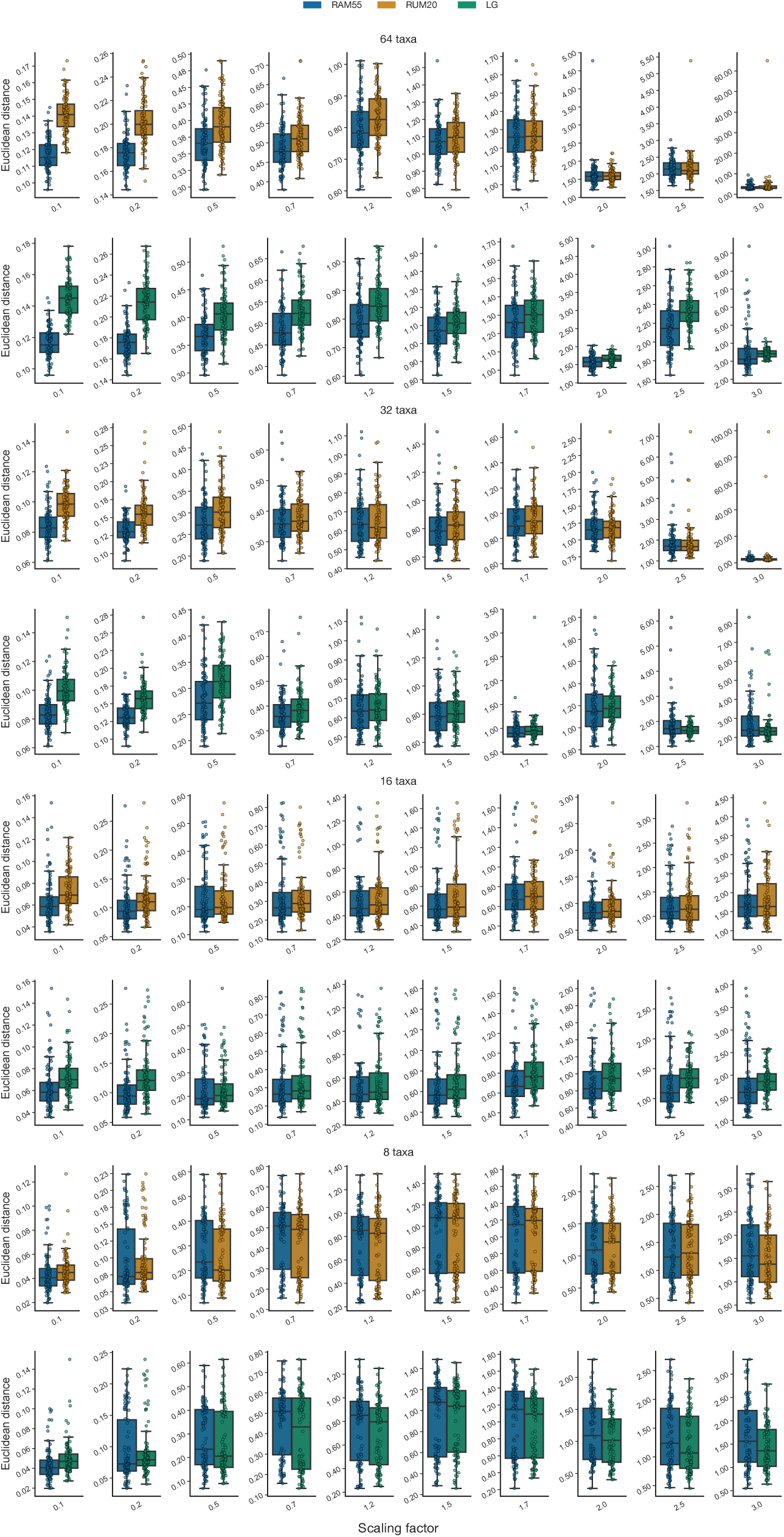
Tree inference accuracy. Rotasequence alignments (200 sites, 100 replicates per scaling) are simulated under RAM55 and the trees in Sup. Fig. 11, scaled according to the factors on the *x*-axis. RUM20, LG and RAM55 itself are then used to perform inference over the simulated rotasequence alignments (or masked amino acid alignments for RAM20 and LG) and the resulting trees are compared to the original phylogeny in terms of Euclidean distance. RAM55 can infer trees that are closer to the original as shown by distance distributions and medians shifted towards lower values.

**Supplementary Figure 4:**
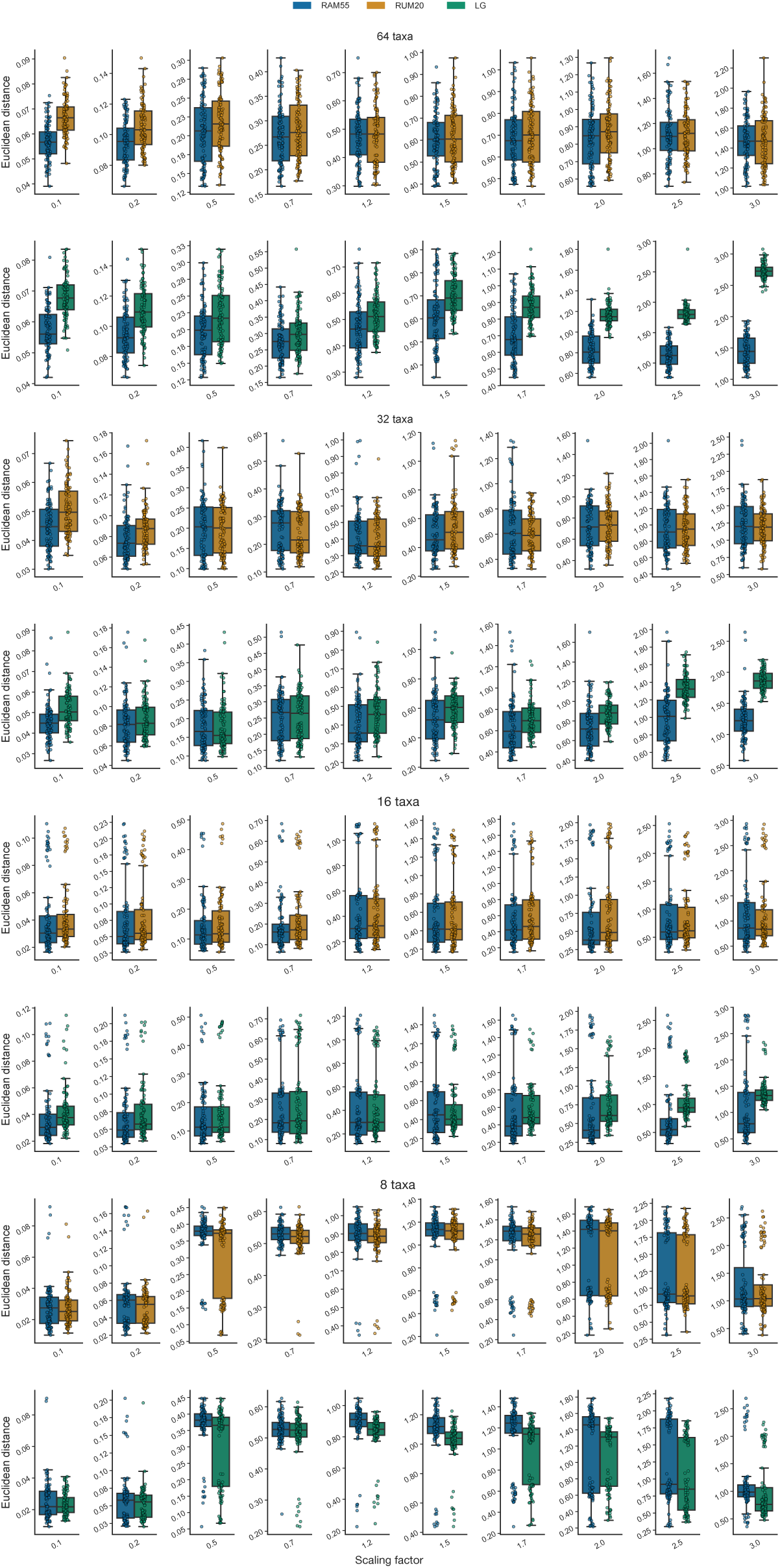
Tree inference accuracy. As Sup. Fig. 3, except simulated alignments each contain 1000 sites.

**Supplementary Figure 5:**
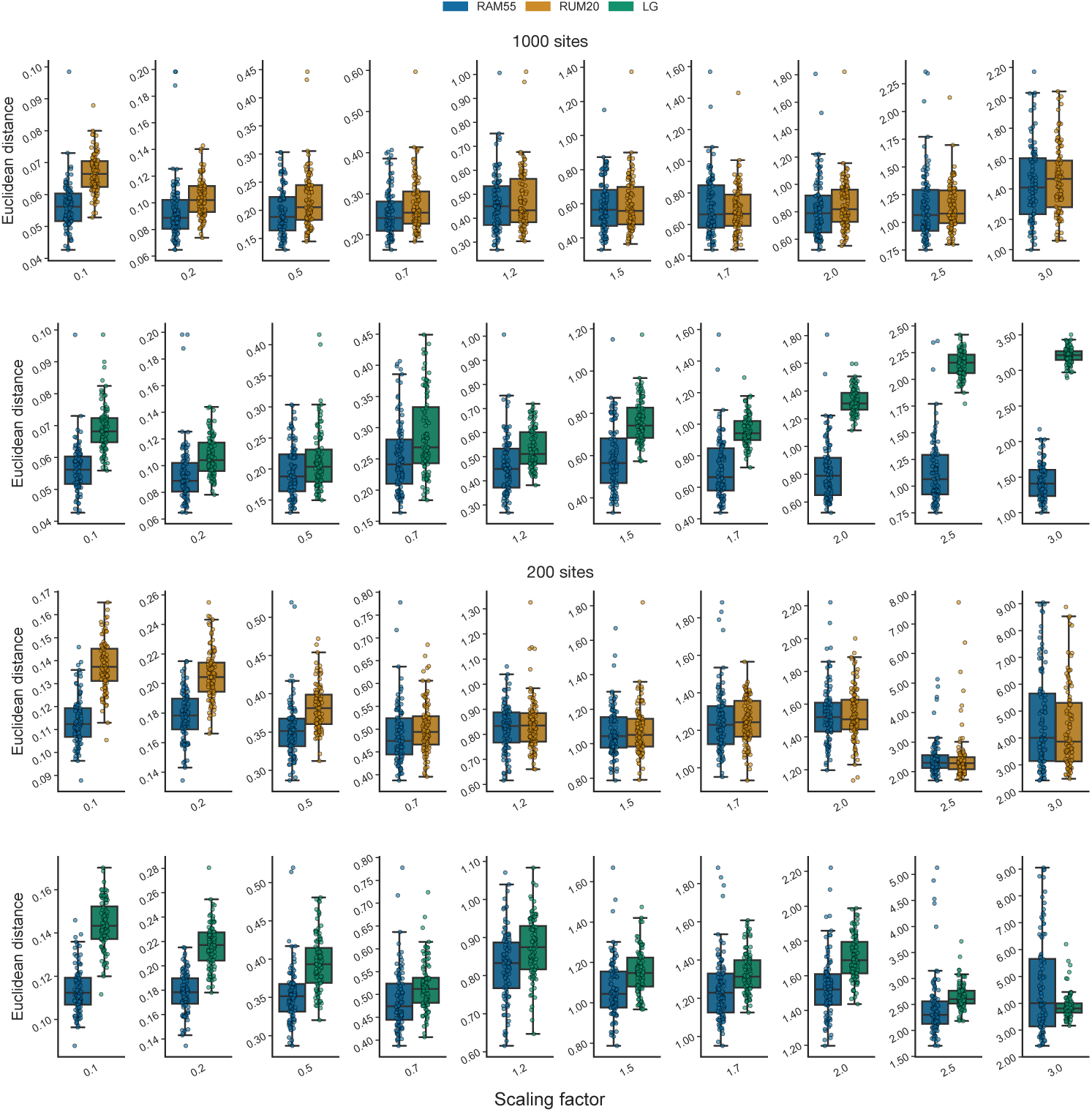
Tree inference accuracy. As Sup. Fig. 3, except simulated alignments are generated under RAM55 and a pruned version of the Ensembl-compara tree (*Tree generation and alignment simulation* and Sup. Files), scaled according to the factors on the *x*-axis.

**Supplementary Figure 6:**
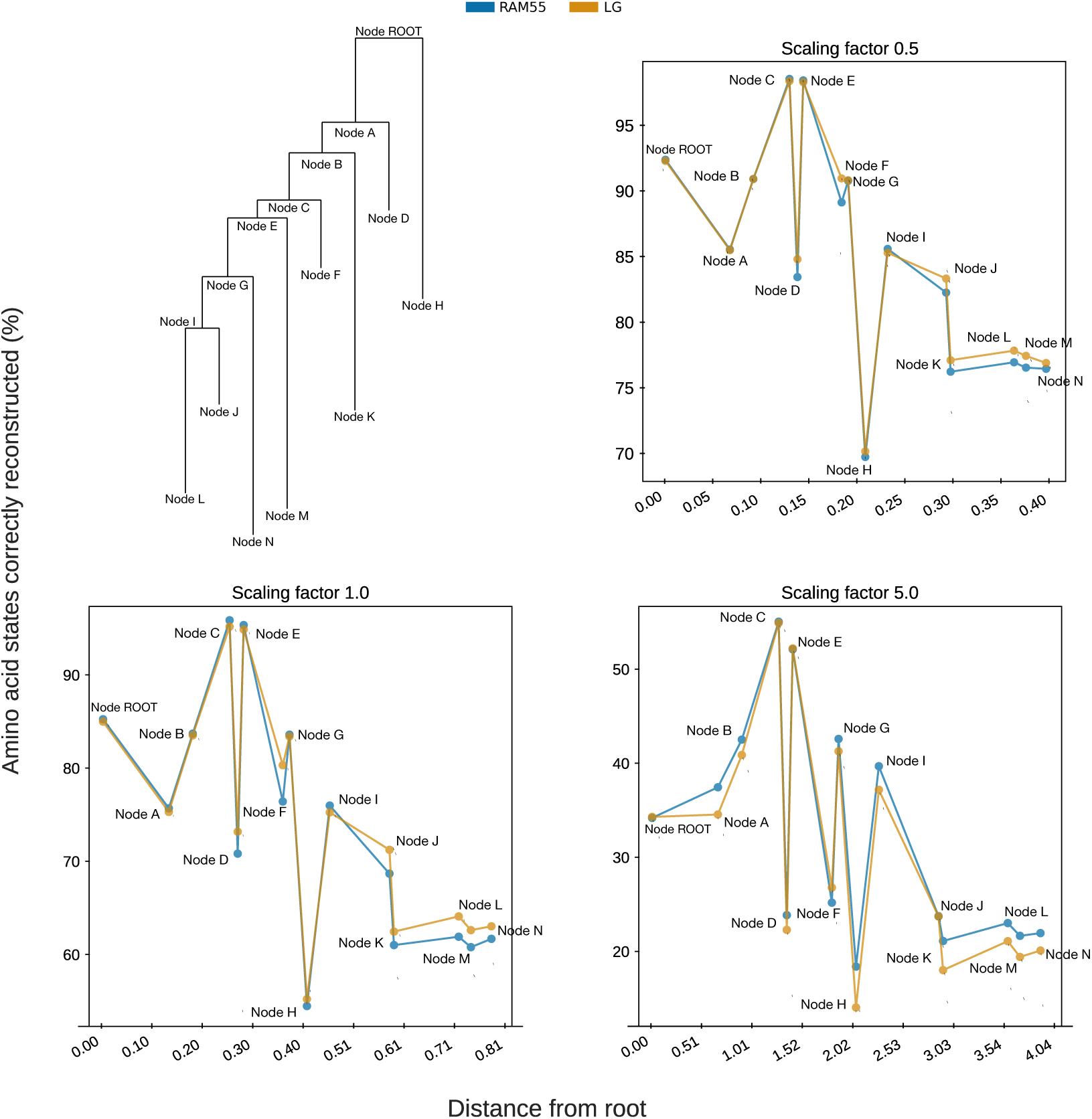
Marginal reconstruction of inferred amino acid accuracy. Amino acid states inferred using marginal reconstruction from rotasequence alignments (200 sites, 8 taxa, 100 replicates per scaling) simulated under RAM55 using our 8-taxon reference phylogeny and scaling its branches according to the factor reported for each sub-plot. The marginal reconstruction algorithm is employed along with RAM55 and our 8-taxa phylogeny to reconstruct internal and terminal nodes’ rotasequences which are then masked to obtain amino acid sequences (in blue, see *Ancestral state reconstruction*). The equivalent procedure is then repeated using LG on ‘masked’ alignments (in orange). Each data point indicates (on the *y*-axis) the mean percentage of amino acid states correctly reconstructed from 100 replicates for a given reconstructed node, against its distance from the root node (*x*-axis).

**Supplementary Figure 7:**
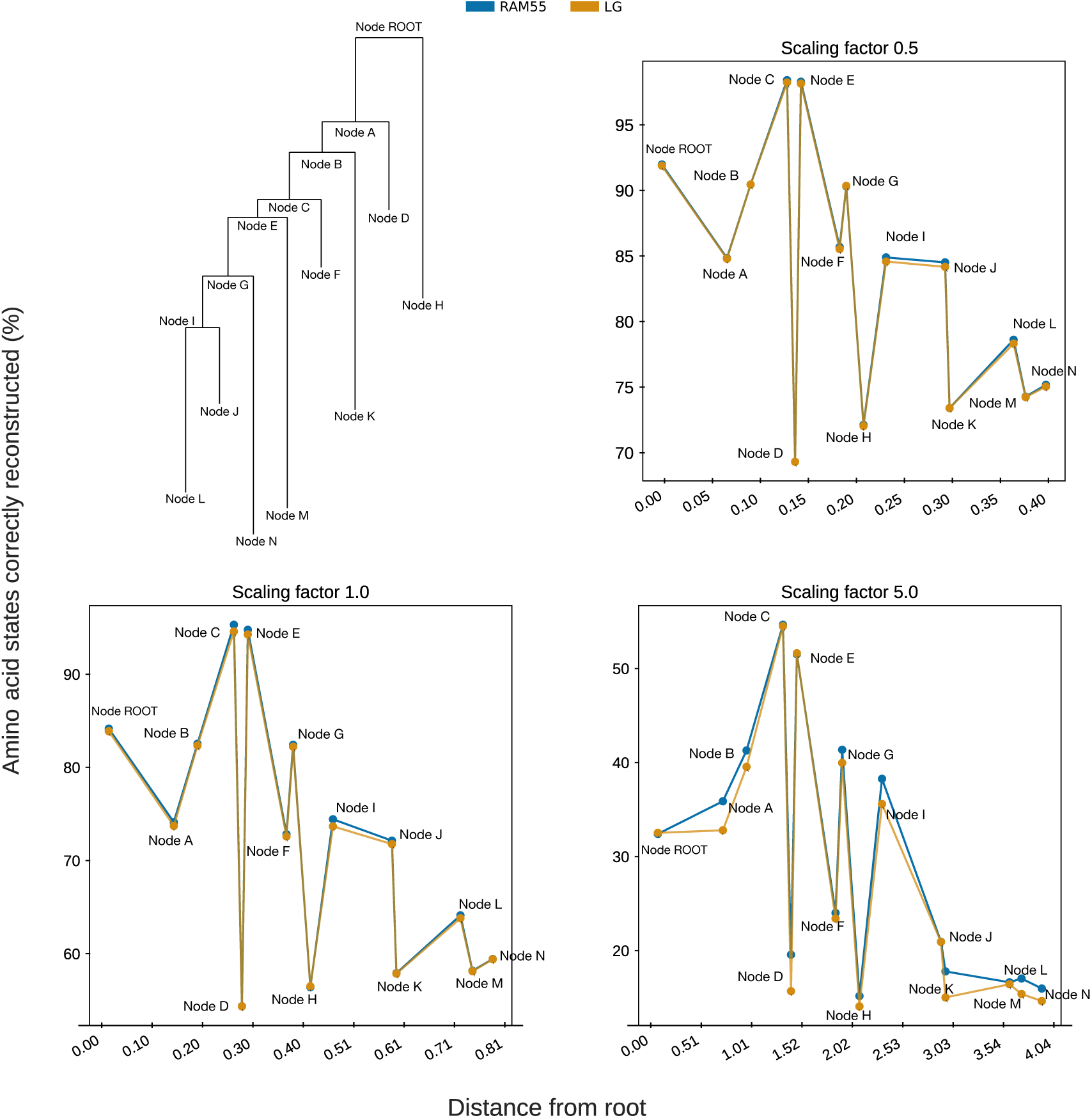
Joint reconstruction of inferred amino acid accuracy. As Sup. Fig. 6, except joint reconstruction is used in place of marginal reconstruction. Marginal and joint reconstruction methods perform about equally in our studies.

**Supplementary Figure 8:**
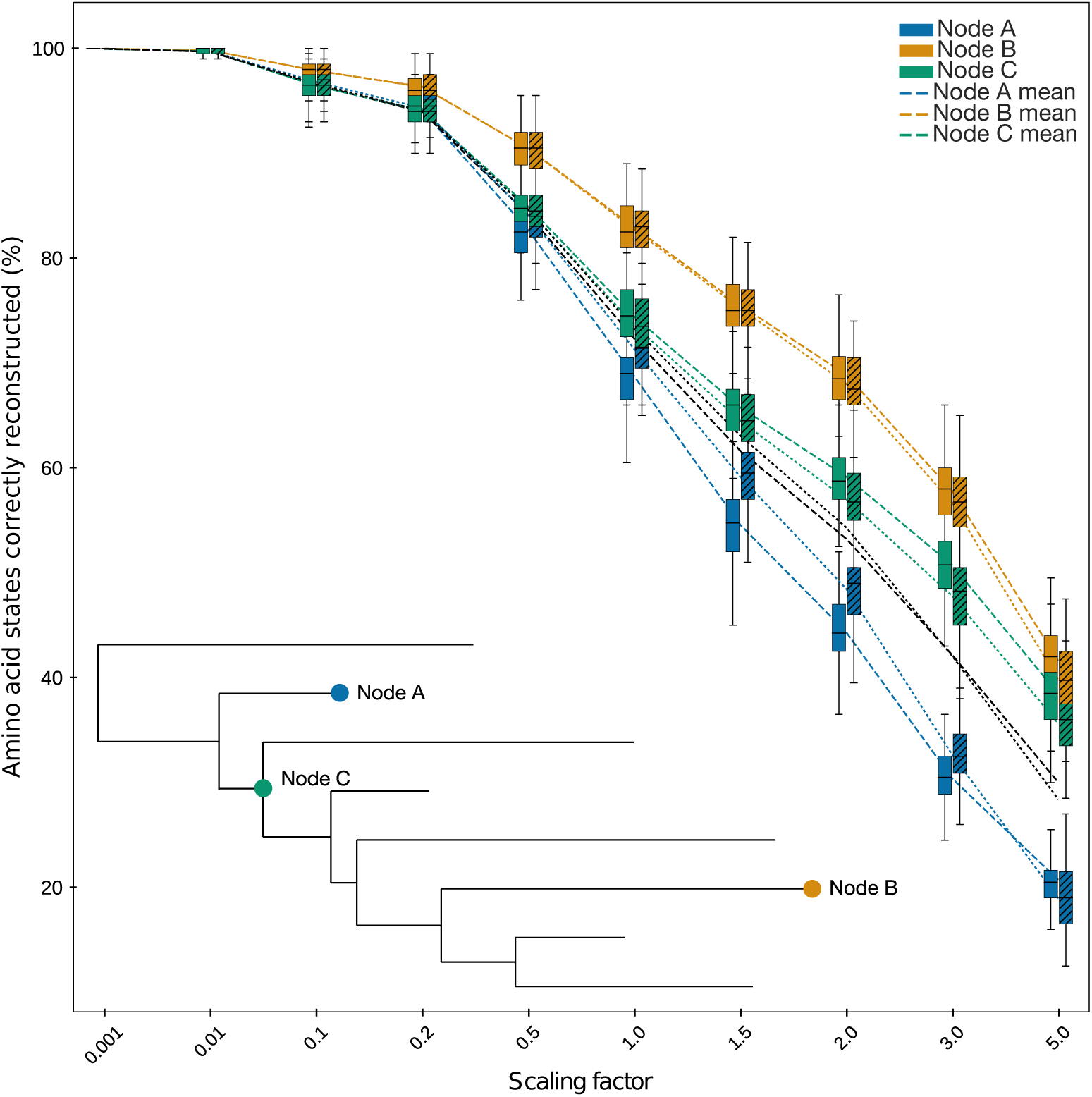
Amino acid state reconstruction accuracy distributions. Ancestral amino acid states are inferred by marginal reconstruction from the same rotasequence alignments as in Fig. 8 using RAM55 and our 8-taxon reference phylogeny, at internal (C) and terminal (A, B) nodes. The same procedure is then repeated using LG on ‘masked’ alignments (hatched). The *y*-axis report the percentage of sites correctly reconstructed for each inferred sequence. Each box-plot contains results from 100 simulation replicates for a given node. Marginal and joint reconstruction methods perform about equally in our studies.

**Supplementary Figure 9:**
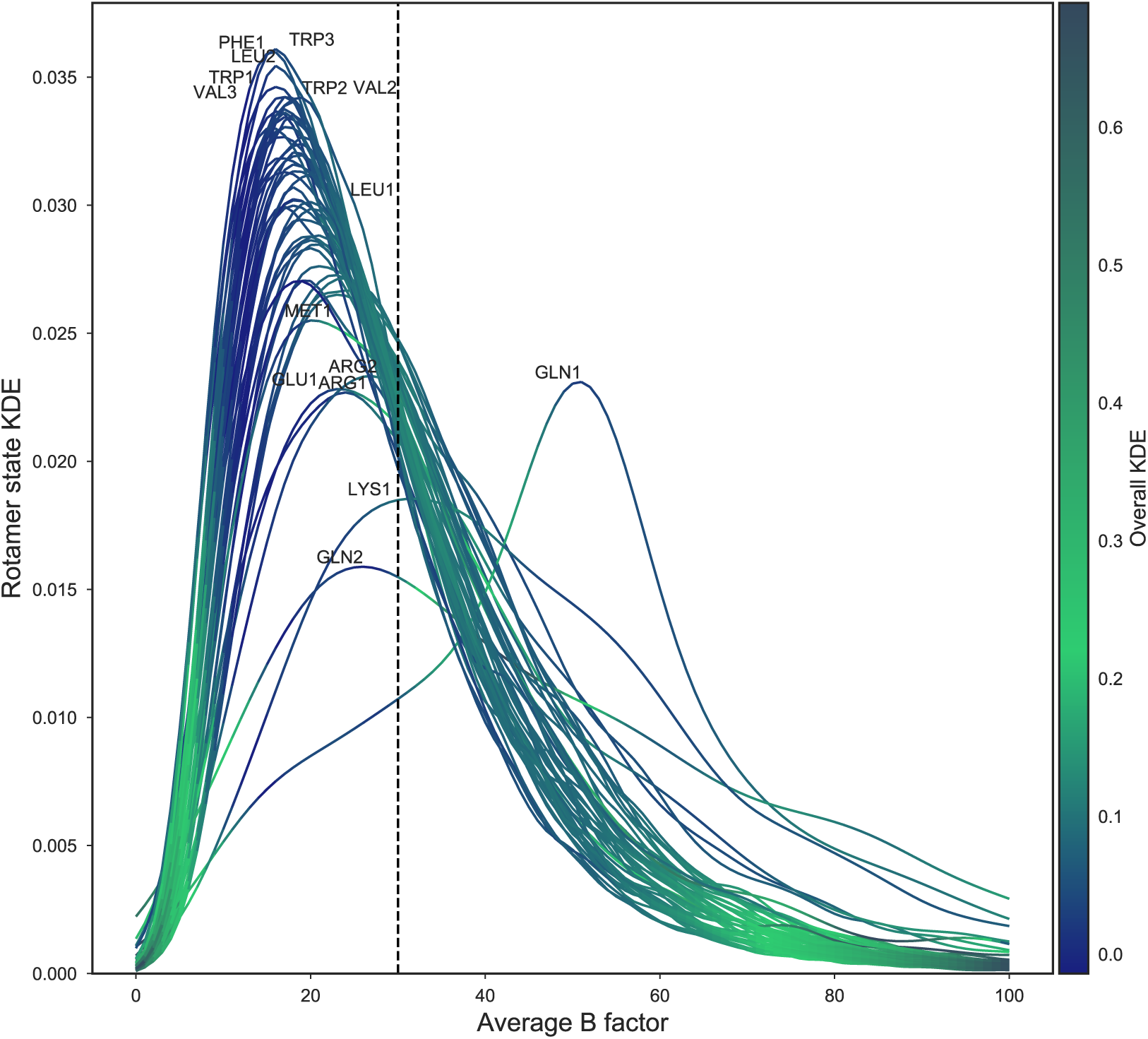
Structural data quality distributions across residues. Kernel density estimate (KDE) for the average B factor for each rotamer state (excluding alanine and glycine) across all residues in our unfiltered dataset. Average B factor is computed over the four atoms (*N, C*^*α*^, *C*^*β*^ and *C*^*γ*^ for most residues) that constitute the dihedral angle defining the *χ*_1_ rotamer configuration. A threshold of B factor < 30 is then applied (dashed line) to ensure only highly reliable atomic coordinates are used to assign rotamer states. Only outlier density plots are labelled, for clarity; the color scheme represents overall plot density distribution at each point along the *x*-axis. Areas under the curves to the right of the dashed line indicates the proportions of each rotamer state in our alignments that are removed by the B factor filtering.

**Supplementary Figure 10:**
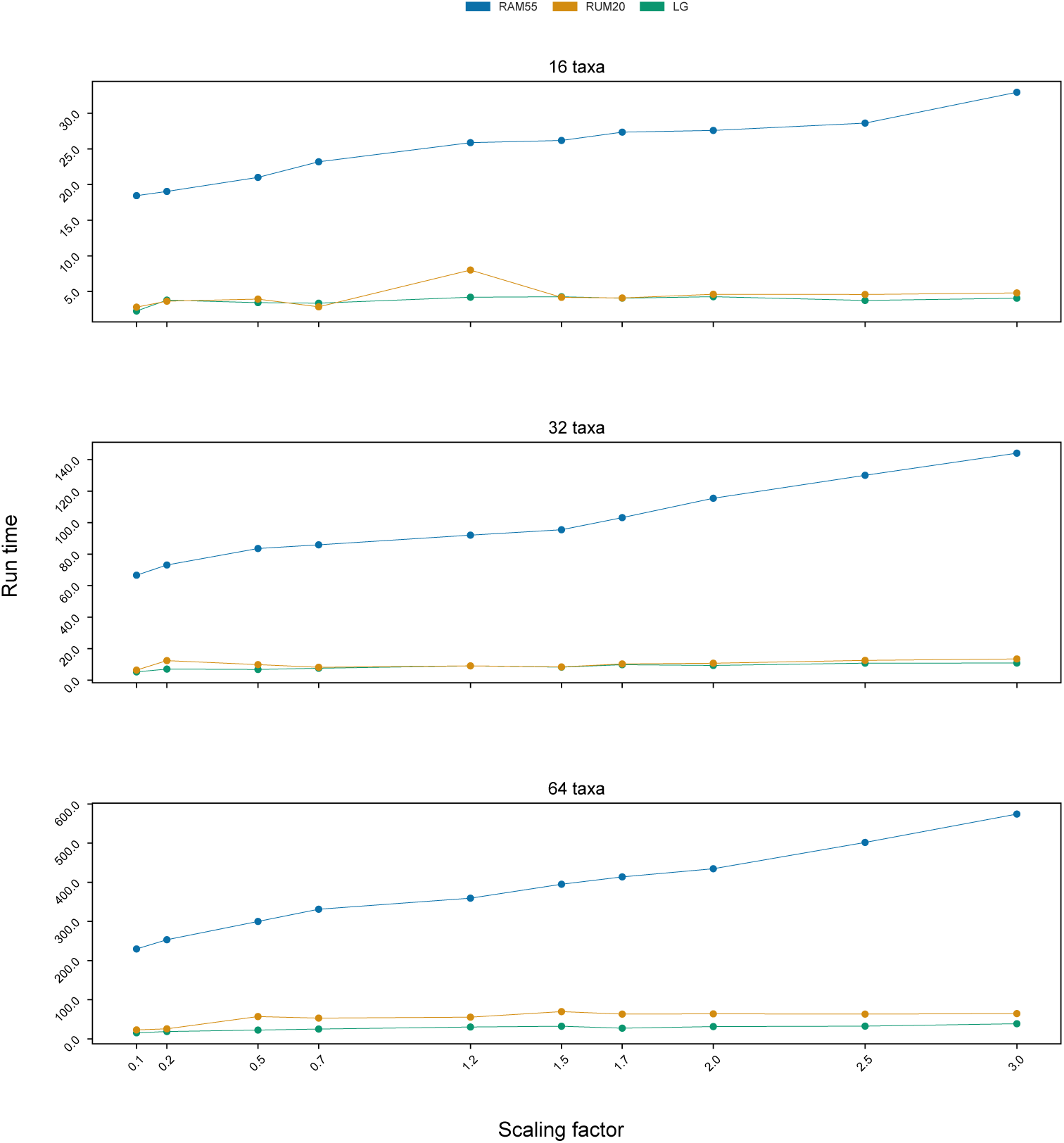
Run time comparisons. Rotasequence alignments (200 sites, 100 replicates per scaling factor) were simulated using RAM55 and the 16-, 32- and 64-taxon trees of Sup. Fig. 11, scaled according to the factors shown on the *x*-axis. The plots report mean run times (in seconds) for ML inference analysis of these rotasequence alignment data under the RUM20, LG and RAM55 models in RAxML-NG (using masked alignments for the RUM20 and LG analyses).

**Supplementary Figure 11:**
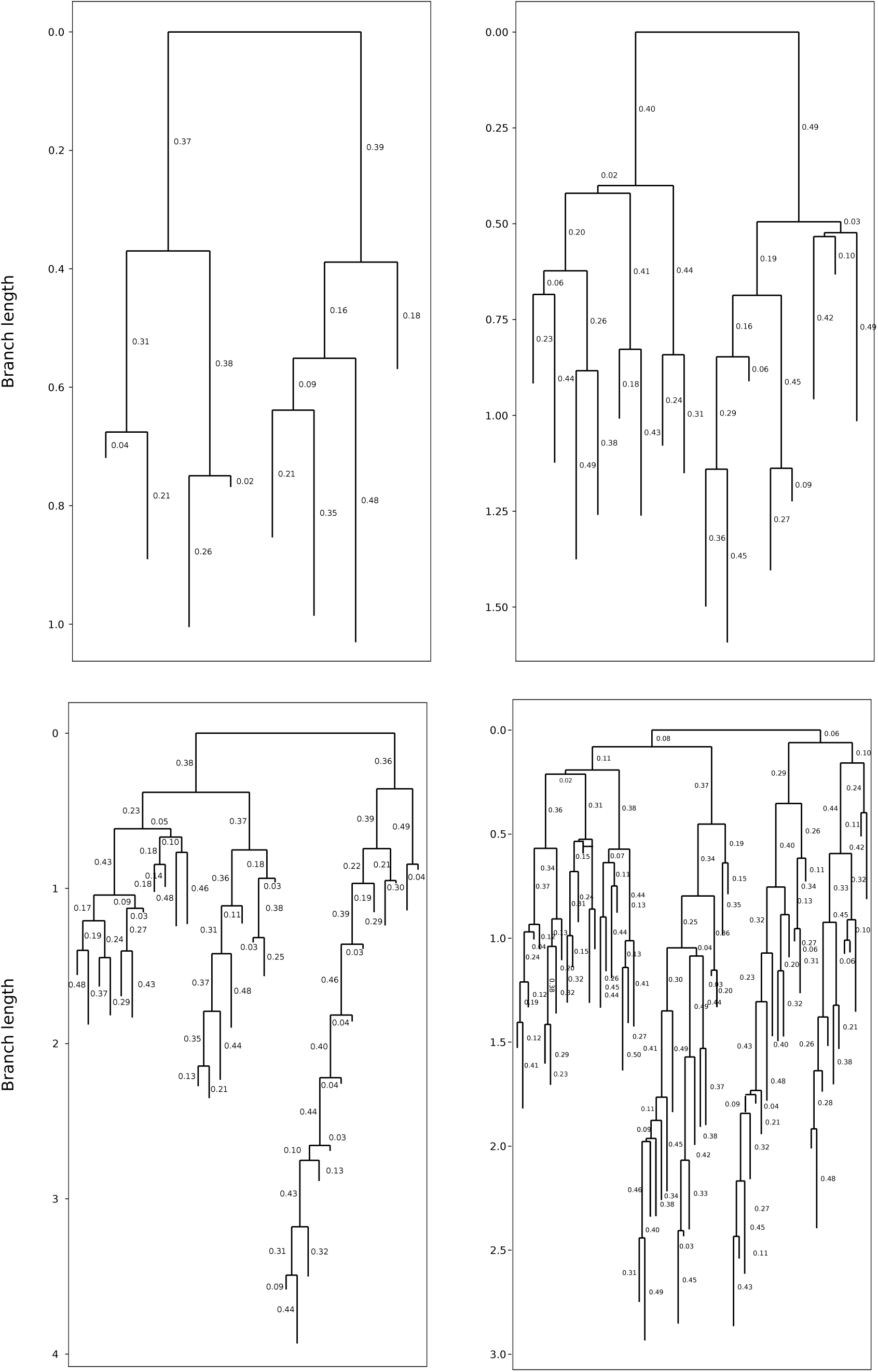
The four randomly-generated phylogenies (8-, 16-, 32- and 64-taxa; available in Sup. Files) used as guides to simulate internal and terminal node sequences (see *Tree generation and alignment simulation, Ancestral state reconstruction*).

**Supplementary Figure 12:**
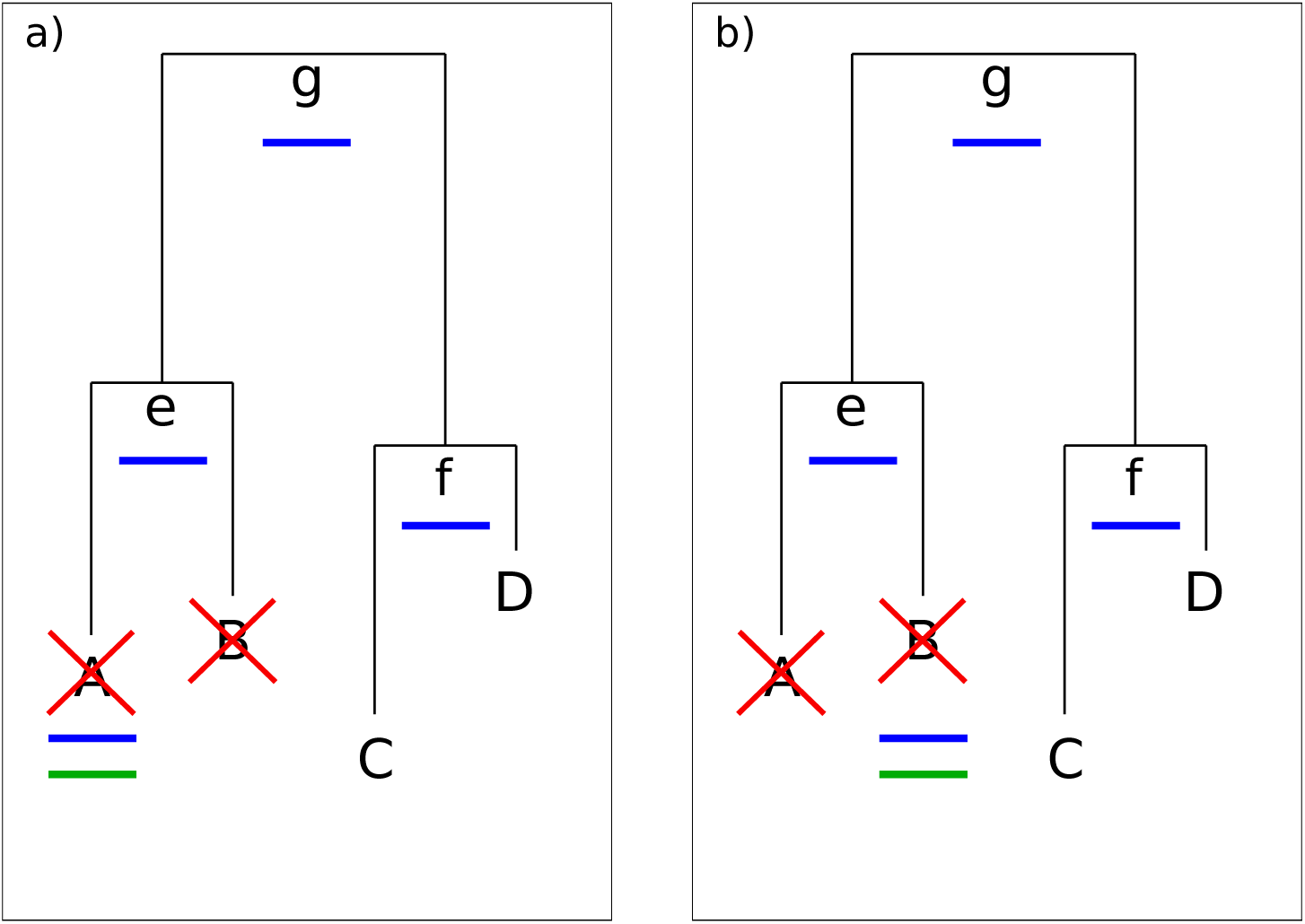
Illustration of the “leave-leaves-out” (LLO) procedure. LLO is used to evaluate a model’s ability to reconstruct terminal nodes’ states. **(a)** A pair of sibling terminal nodes (A, B) is selected, their sequences are removed from the alignment (red ×) and then all internal node sequences are reconstructed (e, f, g, in blue) along with A’s sequence. This can be compared to its original sequence (green). **(b)** The analogous procedure is then followed to reconstruct B’s sequence and compare it to the original.

