## Supplementary material for "Modelling structural constraints on protein evolution via side-chain conformational states": README.txt

The following Supplementary files are included:

all_Pfam_families_tab.tsv	# All Pfam families used to compute RAM55 and RUM20
all_structures_tab.tsv		# All PDB structures used to compute RAM55 and RUM20
PF00514.alignment_rotaseq_1char.PHYLIP	# Empirical rotasequence alignment
Rubisco.alignment_rotaseq_1char.PHYLIP	# Empirical rotasequence alignment
PF07714.alignment_rotaseq_1char.PHYLIP	# Empirical rotasequence alignment
8_leaves.newick	# Randomly-generated berwick tree
64_leaves.newick	# Randomly-generated newick tree
32_leaves.newick	# Randomly-generated newick tree
16_leaves.newick	# Randomly-generated newick tree
RUM20_rates.ng	# RUM20 exchangeabilities for RAxML-NG
RUM20_freqs.ng	# RUM20 amino acid state frequencies for RAxML-NG
RAM55_rates.ng	# RAM55 exchangeabilities for RAxML-NG
RAM55_freqs.ng	# RAM55 rotamer state frequencies for RAxML-NG
RAM55_PAML-style.dat	# Exchangeabilities in PAML-style format. NOTE THAT RAM55 CANNOT BE IMPLEMENTED IN PAML
RUM20_PAML-style.dat	# Exchangeabilities in PAML-style format. NOTE THAT RUM20 CANNOT BE IMPLEMENTED IN PAML
Rubisco.alignment_rotaseq_1char.masked_to_AAseq.PHYLIP	# Empirical rotasequence alignment, masked to an amino acid sequence alignment
PF07714.alignment_rotaseq_1char.masked_to_AAseq.PHYLIP	# Empirical rotasequence alignment, masked to an amino acid sequence alignment
PF00514.alignment_rotaseq_1char.masked_to_AAseq.PHYLIP	# Empirical rotasequence alignment, masked to an amino acid sequence alignment
state_symbols.yaml	# State symbol dictionaries
70_taxa_pruned_ensembl_tree.newick # the Ensembl-compara species tree (https://github.com/Ensembl/ensembl-compara/blob/release/94/scripts/pipeline/species_tree.ensembl.branch_len.nw) pruned down to 70 taxa and scaled so that its average branch length is now 0.257 and thus comparable to our 64-taxa randomly-generated tree (branch length 0.251). These branch lengths, along with the scaling factors used in the simulation process, allow us to cover a realistic range of sequence divergence (~10% to ~85%) while maintaining constant topologies.

Both RAM55 and RUM20 can be implemented using RAxML-NG (https://github.com/amkozlov/raxml-ng).

RAM55 can be specified in RAxML-NG as:
--model MULTI55_GTR{RAM55_rates.ng}+FU{RAM55_freqs.ng}+M{ABCDEFGHIJKLMNOPQRSTUVWYXZabcdefghijklmnopqrstuvwxyz012}{-}

RUM20 can be specified in RAxML-NG as:
--model MULTI20_GTR{RUM20_rates.ng}+FU{RUM20_freqs.ng}+M{ARNDCQEGHILKMFPSTWYV}{-}
