## Supplementary material for "Modelling structural constraints on protein evolution via side-chain conformational states": state_symbols.yaml

```
rotamer_one2four_dict:
  '0': VAL1
  '2': VAL3
  B: ARG1
  D: ARG3
  F: ASN2
  H: ASP1
  J: ASP3
  L: CYS2
  N: GLN1
  P: GLN3
  R: GLU2
  T: GLY
  V: HIS2
  X: ILE1
  Z: ILE3
  b: LEU2
  d: LYS1
  f: LYS3
  h: MET2
  j: PHE1
  l: PHE3
  n: PRO2
  p: SER2
  r: THR1
  t: THR3
  v: TRP2
  x: TYR1
  z: TYR3
  '1': VAL2
  A: ALA
  C: ARG2
  E: ASN1
  G: ASN3
  I: ASP2
  K: CYS1
  M: CYS3
  O: GLN2
  Q: GLU1
  S: GLU3
  U: HIS1
  W: HIS3
  Y: ILE2
  a: LEU1
  c: LEU3
  e: LYS2
  g: MET1
  i: MET3
  k: PHE2
  m: PRO1
  o: SER1
  q: SER3
  s: THR2
  u: TRP1
  w: TRP3
  y: TYR2
AA_one2three_dict, rotamer_masking_dict:
  A: ALA
  C: CYS
  E: GLU
  D: ASP
  G: GLY
  F: PHE
  I: ILE
  H: HIS
  K: LYS
  M: MET
  L: LEU
  N: ASN
  Q: GLN
  P: PRO
  S: SER
  R: ARG
  T: THR
  W: TRP
  V: VAL
  Y: TYR
rotamer_four2one_dict:
  TYR3: z
  TYR2: y
  TYR1: x
  ASP2: I
  HIS3: W
  ASN2: F
  PRO2: n
  PRO1: m
  ASN1: E
  GLY: T
  GLU1: Q
  CYS1: K
  CYS3: M
  CYS2: L
  TRP3: w
  PHE3: l
  PHE2: k
  PHE1: j
  ILE1: X
  ILE2: Y
  ILE3: Z
  VAL3: '2'
  VAL2: '1'
  VAL1: '0'
  TRP1: u
  SER2: p
  SER3: q
  SER1: o
  ASN3: G
  TRP2: v
  LEU2: b
  ASP1: H
  HIS2: V
  ASP3: J
  LYS1: d
  LYS2: e
  LYS3: f
  THR2: s
  THR3: t
  THR1: r
  MET2: h
  MET3: i
  MET1: g
  LEU3: c
  ARG1: B
  ARG2: C
  ARG3: D
  GLU2: R
  GLU3: S
  ALA: A
  LEU1: a
  GLN3: P
  GLN2: O
  GLN1: N
  HIS1: U
AA_three2one_dict:
  CYS: C
  ASP: D
  SER: S
  GLN: Q
  LYS: K
  ASN: N
  PRO: P
  THR: T
  PHE: F
  ALA: A
  HIS: H
  GLY: G
  ILE: I
  LEU: L
  ARG: R
  TRP: W
  VAL: V
  GLU: E
  TYR: Y
  MET: M
```
